# Osmotic niche changes as multifaceted trigger of cellular regenerative processes in organ injury

**DOI:** 10.1101/2025.09.07.674644

**Authors:** Matthias R. Kollert, Taimoor H. Qazi, Mai H.Q. Phan, Blanka Majchrzycka, Raphael S. Knecht, Tobias Thiele, Denise Jahn, Raimund Schlüßler, Andreas F. Thünemann, Peter Fratzl, Daniel M. Ibrahim, Viola Vogel, David J. Mooney, Georg N. Duda

## Abstract

Tissue compartments can maintain distinct osmotic environments that are separated by physical barriers. Whether injury-induced osmotic niche changes associated with the disruption of these barriers contribute to the onset of regeneration remains unknown. Here we present analyses of human tissue samples indicating that fracture-induced barrier disruption likely exposes the marrow niche to a sudden hypo-osmotic shift, which enhances extracellular matrix (ECM) stress relaxation. We reveal that this hypo-osmolar shift, mimicked in vitro by lowering ion concentrations, triggers bone regenerative processes in mesenchymal stromal cells (MSCs), markedly increasing their spreading, proliferation, and osteogenic differentiation while residing in osmolality-responsive viscoelastic ECM. In elastic ECM, elevated ion concentrations instead promoted osteogenic differentiation, indicating that ECM viscous dissipation redirects how cells interpret osmotic changes. Mechanistically, the regenerative function of the osmolality drop depends on the matrix providing cell-adhesion ligands and physiological viscoelastic properties. Within eight hours, hypo-osmolality amplifies pro-inflammatory and pro-angiogenic cytokine secretion (CXCL1, CCL2, CCL5, and IL-6). Within two hours, transcriptomic analyses reveal chromatin remodeling signatures specifically associated with hypo-osmolality. Within minutes, the osmolality drop alters nuclear morphology, mechanics, and chromatin organization, preceding increased osteogenic transcription factor expression. These findings identify osmotic niche disruption as a fast-acting physical signal that coordinates ECM mechanics and nuclear organization to initiate regenerative programs after injury. This osmotic stimulus may coordinate regenerative responses beyond the stromal compartment through inflammatory signaling. Leveraging osmotic cues to induce regenerative cascades may eventually advance local and personalized therapeutic strategies for patients with impaired healing capacity. The integration of osmotic and mechanical ECM properties, as demonstrated in our assay, may catalyze advanced 3D cell culture systems and offer new perspectives on material design in tissue engineering, disease modeling, and mechanobiology.

## Main

Living tissues maintain distinct osmotic environments between compartments separated by barriers that restrict solute and water exchange. Osmotic gradients are central to physiology and tissue mechanics, for example in kidney function^1^ and load-bearing connective tissues^2,3^, and can be dysregulated in pathology, including inflammatory swelling and edema formation^4^ or cancer-associated osmotic differences^5,6^. Whether abrupt disruption of compartment boundaries during tissue injury produces osmotic niche changes that contribute to the initiation of regeneration remains unclear.

The bone marrow niche resides within a continuously remodeled mineralized tissue, suggesting an extracellular environment whose ionic and pH conditions differ from those of surrounding soft tissues and fluids^7,8^. Consistent with this, our osmolality measurements of ex vivo human tissue samples showed that bone marrow exhibits higher osmolality than peripheral blood and cerebrospinal fluid (Fig. 1a). This suggests that fracture of the cortical bone barrier likely exposes the marrow niche to a sudden osmolality drop, an event that coincides with, and may contribute to, the initiation of regeneration.

**Fig. 1.**
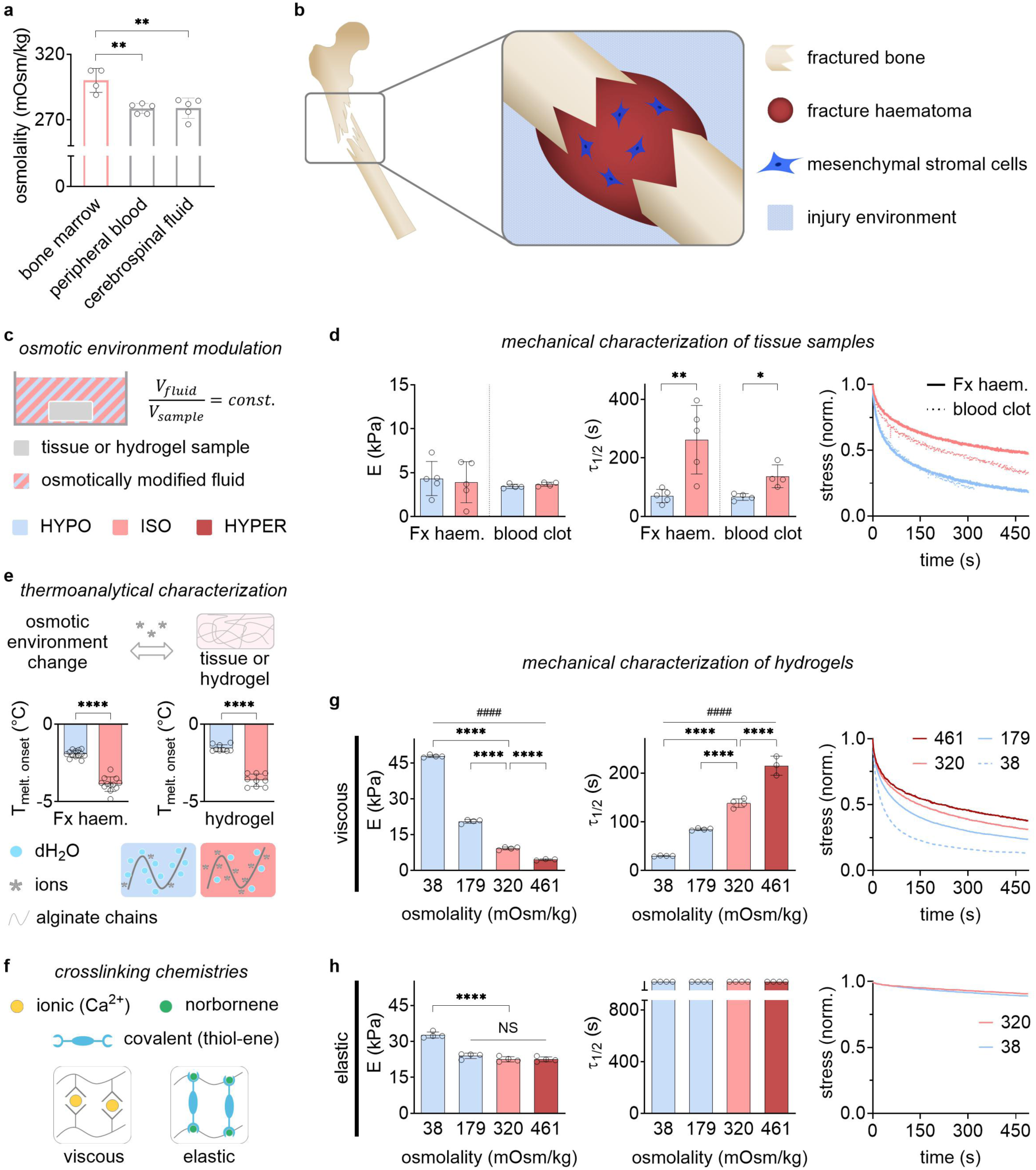
Environmental osmolality regulates the mechanics of fracture haematoma and hydrogels. (**a**) Osmolality of ex vivo human samples of the bone marrow niche and of tissues typically surrounding bone in the event of fracture (i.e., peripheral blood, cerebrospinal fluid) (*n* = 4–5 donors, \*\**P* < 0.01 by one-way ANOVA). (**b**) Illustration of the injury site immediately after bone fracture and disruption of the bone marrow niche, leading to exposure of fracture haematoma with residing cells (e.g., mesenchymal stromal cells) to altered environmental properties (e.g., lower osmolality). (**c**) Experimental setup to modulate the osmotic environment of a tissue sample or hydrogel. After incubation of a tissue sample or hydrogel in hypo-osmolar (HYPO) or iso-osmolar (ISO) environment, mechanical characterization was performed. Elastic and viscous (i.e., stress relaxing) mechanical properties were quantified by elastic modulus, *E*, and stress relaxation half time, *τ*_1⁄2_, respectively. (**d**) *E* (left), *τ*_1⁄2_ (middle) and representative stress relaxation curves (right) of ex vivo human fracture haematoma (Fx haem.) and in vitro coagulated human blood clots (*n* = 5 donors of Fx haem., *n* = 4 samples of in vitro blood clots, \*\**P* < 0.01, and \**P* < 0.05 by Student’s t test). (**e**) Thermoanalytical characterization was performed using DSC on tissue samples (Fx haem.) and alginate hydrogels after incubation in HYPO or ISO solutions. Changes in melting onset temperature (T_melt_) would indicate alterations in material properties. Top: Illustration of how altered environmental osmolality (i.e., ion concentration) may affect tissue or hydrogel properties due to ion diffusion; middle: melting onset temperature (T_melt_) of fracture haematoma samples or alginate hydrogels after osmotic modulation suggest increased water binding strength due to ion efflux from material samples in HYPO environments (Fx haem.: *n* = 12–13 samples from 5 donors, hydrogel: *n* = 9 samples, \*\*\*\**P* < 0.0001 by Student’s t test); bottom: illustration of increased water on alginate polymer chains after ion efflux in HYPO environments. (**f**) Illustrations of ionic (left) and thiol-ene covalent click (right) chemistries used for alginate crosslinking to produce viscoelastic hydrogels with pronounced stress relaxation properties (viscous) and predominantly elastic hydrogels (elastic), respectively. (**g**) Viscous hydrogels exhibited an osmo-mechanical interplay similar to that identified in fracture hematomas and blood clots, characterized by faster stress relaxation with decreasing buffer concentration (*n* = 4 samples, \*\*\*\**P* < 0.0001 by one-way ANOVA, comparisons to ISO are shown, ####*P* < 0.0001 by Spearman’s rank correlation). (**h**) Elastic hydrogels with elastic mechanical properties which do not exhibit a similar osmo-mechanical interplay (*n* = 4 samples, \*\*\*\**P* < 0.0001, NS: not significant (*P* > 0.05) by one-way ANOVA, comparisons with ISO are shown). Bar plots: mean ± standard deviation (SD).

Changes in environmental ion concentration can alter extracellular matrix (ECM) mechanics through charge-mediated interactions with negatively charged ECM components^3,9,10^, and influence cellular ion homeostasis through regulatory volume adaptation^11,12^. Even moderate shifts in ionic strength can substantially alter swelling pressure and elastic properties in charged ECM by tens of kPa^3,9,13^, a magnitude of mechanical change sufficient to influence cell fate^14^. These ion-driven osmotic changes are mechanistically distinct from osmotic pressure exerted by macromolecules such as polyethylene glycol (PEG), which persistently constrict cell volume^11,15^ and dehydrate tissue^16–18^. How cells respond to osmotic changes is influenced by cytoskeleton organization^11,12^, which in turn is shaped by ECM mechanics^19,20^.

Cells can sense ECM mechanical properties, for example through integrin-mediated adhesion to the ECM, translating physical cues into biochemical signals^20–23^. ECM viscoelasticity, and specifically stress relaxation, has emerged as a key regulator of cell fate, driving essential bone regenerative processes including spreading, proliferation, and osteogenic differentiation of mesenchymal stromal cells (MSCs) ^14,24^. Bone fracture healing is widely considered a model of scar-free regeneration after injury^25^. Particularly the early post-fracture phase^25,26^, characterized by fracture haematoma with viscoelastic ECM properties^14^, is a critical determinant of healing outcome. However, whether the osmolality drop that likely accompanies fracture constitutes a regenerative signal for resident cells through integrated osmotic-mechanical changes remains unexplored.

We postulated that the bone marrow niche, where MSCs reside, experiences an acute osmolality drop upon bone fracture (Fig. 1b). This hypo-osmolar niche change likely results from altered concentrations of small osmolytes such as ions, affects both the ECM and resident cells, and occurs prior to the onset of regeneration. To dissect how these coupled extracellular and intracellular effects influence early regenerative processes, we developed an in vitro platform combining osmotically tunable viscoelastic hydrogels with 3D cell culture, which enabled systematic modulation of niche osmolality while controlling ECM mechanics and cell-matrix adhesion.

### Lowered ion concentration enhances fracture haematoma stress relaxation

To investigate how osmotic changes influence ECM mechanics, we established an experimental setup that allows to modulate the osmotic environment of tissue samples or hydrogels (Fig. 1c). The setup allowed for deliberate tuning of the osmotic environment of a tissue, or hydrogel, sample by adjusting the osmotic concentration of the fluid (i.e., buffer) while maintaining a constant fluid-to-sample volume ratio under controlled incubation conditions. First, we aimed to understand how a drop in osmolality would affect the viscoelastic properties of the ECM surrounding MSCs (i.e., bone marrow). We incubated freshly dissected ex vivo samples of human initial fracture haematoma in iso-osmolar (ISO) or extremely hypo-osmolar (HYPO) environments established using PBS or ddH_2_O. The measured elastic modulus (*E*) and stress relaxation half-time (*τ*_1⁄2_) of fracture haematoma in ISO condition corresponded to previously reported values for the tissue in fresh condition^27,28^ (Fig. 1d, Supplementary Fig. 1). This suggested that the mechanical properties of samples in ISO condition were similar to native bone marrow. Incubation in HYPO condition did not significantly alter *E*, but clearly decreased *τ*_1⁄2_, indicating faster stress relaxation, or more viscous, material characteristics. For validation, we characterized in vitro produced (human) blood clots, which exhibited lower *τ*_1⁄2_ in HYPO compared to ISO conditions, consistent with the results for the fracture haematomas. These results agree with previous research showing that incubation of ex vivo tendon samples in hypo-osmolar conditions, established by lowered ion buffer concentration, led to faster ECM stress relaxation^10^. These findings are remarkable as faster ECM stress relaxation was previously shown to promote regenerative processes of MSCs, such as osteogenic differentiation^14^. Our data now suggest that the drop in osmolality, which the niche of MSCs may experience during bone fracture, enhances the stress relaxation properties of the (bone marrow) ECM of MSCs. Thus, a sudden change in niche osmolality might serve as a trigger of MSC behavior through tuning ECM mechanics.

Modulating buffer ion concentration can influence the mechanics (*E*) of tissue samples by altering electrostatic effects, as reported in cartilage^3,9^. Keeping in mind that surface charge effects are related to water binding properties, we used differential scanning calorimetry (DSC). DSC can detect potential differences in the melting onset temperature (T_melt_) between incubated material samples, which could indicate differences in water binding properties^29,30^. T_melt_ of fracture haematoma samples was indeed higher in HYPO compared to ISO conditions (Fig. 1e), suggesting that higher charge effects may have caused the enhanced ECM viscoelasticity in HYPO conditions (Fig. 1d).

### Hydrogels to mimic the interplay between osmolality and ECM stress relaxation

We sought to emulate the interplay between environmental osmolality and ECM stress relaxation properties in a controlled manner using synthetic biomaterials that can serve as artificial ECM for in vitro cell culture. We used ionically crosslinked alginate hydrogels for 3D cell culture, as these have been shown previously to be suitable for mimicking viscoelastic ECM properties in vitro^14^. DSC characterization of these hydrogels showed higher T_melt_ in HYPO (ddH_2_O) compared to ISO (DMEM) conditions, suggesting comparable changes in water binding, and charge, properties as observed in fracture haematoma samples (Fig. 1e). To better understand this effect, we performed more detailed characterization of the synthetic (hydrogel) material, as its properties are more reproducible compared to ex vivo tissue samples (Fx haem.). Freezable water content (W_f_), swelling, and dry weight were quantified (Supplementary Fig. 2). Measurements of hydrogels in HYPO compared to ISO conditions showed a higher W_f_, almost no swelling, and a lower dry weight. This suggested that altered water binding properties were likely caused by ion efflux from the hydrogels, indicating elevated charge effects in the hydrogels in HYPO compared to ISO conditions.

Next, we established two distinct hydrogel systems, suitable for 3D cell culture, in our experimental setup to decouple the effects of osmotic changes on the ECM mechanics from those on the encapsulated cells. The two hydrogel systems were designed to investigate: i) the role of the mechanical characteristics of the ECM (i.e., viscoelastic versus predominantly elastic), and ii) the role of the identified interplay between osmolality and ECM stress relaxation on cell regenerative processes. In experiments assessing the tuneability of hydrogel mechanics by osmotic environment modulation, the osmotic concentration of the buffer (DMEM) was adjusted through dilution with ddH_2_O. For hydrogels with more physiological viscoelastic properties, we used ionic (calcium-mediated) crosslinking (Fig. 1f, left). Strikingly, these hydrogels exhibited a relationship between osmolality and ECM stress relaxation similar to that observed in ex vivo human fracture haematomas: lower environmental osmolality resulted in enhanced ECM stress relaxation (Fig. 1g, Fig. 1d). To assess whether calcium-mediated crosslinking of the hydrogel was altered in HYPO compared with ISO conditions, calcium concentration in hydrogels was measured by ICP-MS after one day of incubation. No significant differences in calcium content were detected in the tested osmolality range (Supplementary Fig. 3). Together with previous work showing that hypo-osmolar environments enhance ECM stress relaxation through glycosaminoglycan charge effects in ex vivo tendons^10^, our data suggest that, under HYPO conditions, increased electrostatic repulsion between anionic alginate polysaccharide chains may enhance hydrogel stress relaxation. Our data show that the viscoelasticity of calcium-crosslinked alginate hydrogels can be tuned by modulating their osmotic environment (i.e., ion concentration), or the volume ratio between surrounding fluid and hydrogel (Supplementary Fig. 4). To contrast the observed mechanical characteristics (i.e., interplay between osmolality and stress relaxation; viscoelasticity of ECM), we used thiol-ene click chemistry to covalently crosslink norbornene-modified alginate chains to limit stress relaxation (Fig. 1f, right), and high molecular weight alginate to decrease the influence of altered buffer ionic strength^31,32^. The hydrogels obtained exhibited predominantly elastic material properties and showed a moderate change in *E* only in the extremely hypo-osmolar condition, but remained largely unaffected in a wide osmolality range (179 to 461 mOsm/kg) (Fig. 1h). Looking into the effects of ECM covalent crosslinking is sign, since the native ECM is crosslinked and partially cleaved enzymatically in a wound site. Due to the contrasting values for *τ*_1⁄2_, the ionically crosslinked hydrogels with also liquid material characteristics were termed as ‘viscous’, and the non-relaxing covalently crosslinked hydrogels as ‘elastic’.

### Osmotic changes regulate MSC spreading and proliferation in viscous, but not in elastic hydrogels

To investigate how acute changes in niche osmolality may affect the onset of regeneration, we studied the early phase of osteogenic differentiation of MSCs. The osmolality drop in the marrow niche upon bone fracture would likely be rooted in the barrier function of the cortical bone cavity, which separates the marrow from the lower osmolality in tissues surrounding bone (Fig. 2a). An acutely lowered osmolality in the injured niche would likely prevail for several days after fracture, because the cortical bone barrier is not repaired within seven days post injury (DPI) even in mice^33^. The impact of osmotic changes in osteogenic differentiation of MSCs was investigated using an established induction protocol^14^. The cells were encapsulated and equilibrated in 3D culture with the mechanically defined hydrogel systems presented above and with integrated osmotic environment modulation through modified incubation medium (Fig. 2b). Considering the extended culture durations, we aimed to avoid confounding effects of extreme osmotic conditions (e.g., stress, apoptosis) by defining an osmolality range maintaining cell viability and metabolic activity comparable to standard culture conditions. We defined a formulation that modified medium osmolality via ion buffer concentration, while maintaining consistent concentrations of medium supplements (e.g., serum, differentiation-inducing factors) (Supplementary Fig. 5). With this approach, a focus osmolality range of 250–390 mOsm/kg was established that allowed to model acute osmotic niche changes in hydrogel-based 3D culture. For the following experiments, 250, 320, and 390 mOsm/kg served as reference values of osmolality in HYPO, ISO, and HYPER conditions, respectively. Accordingly, the osmolality drop modelled in vitro represented a ∼22% change, and thus was of similar order of magnitude as the osmolality drop of ∼8% measured in ex vivo fluids (Fig. 1b). Previous studies have shown that osmolality changes in this range can have substantial impact in biological tissues, for example in swelling pressure and ECM mechanics^13,16^.

**Fig. 2.**
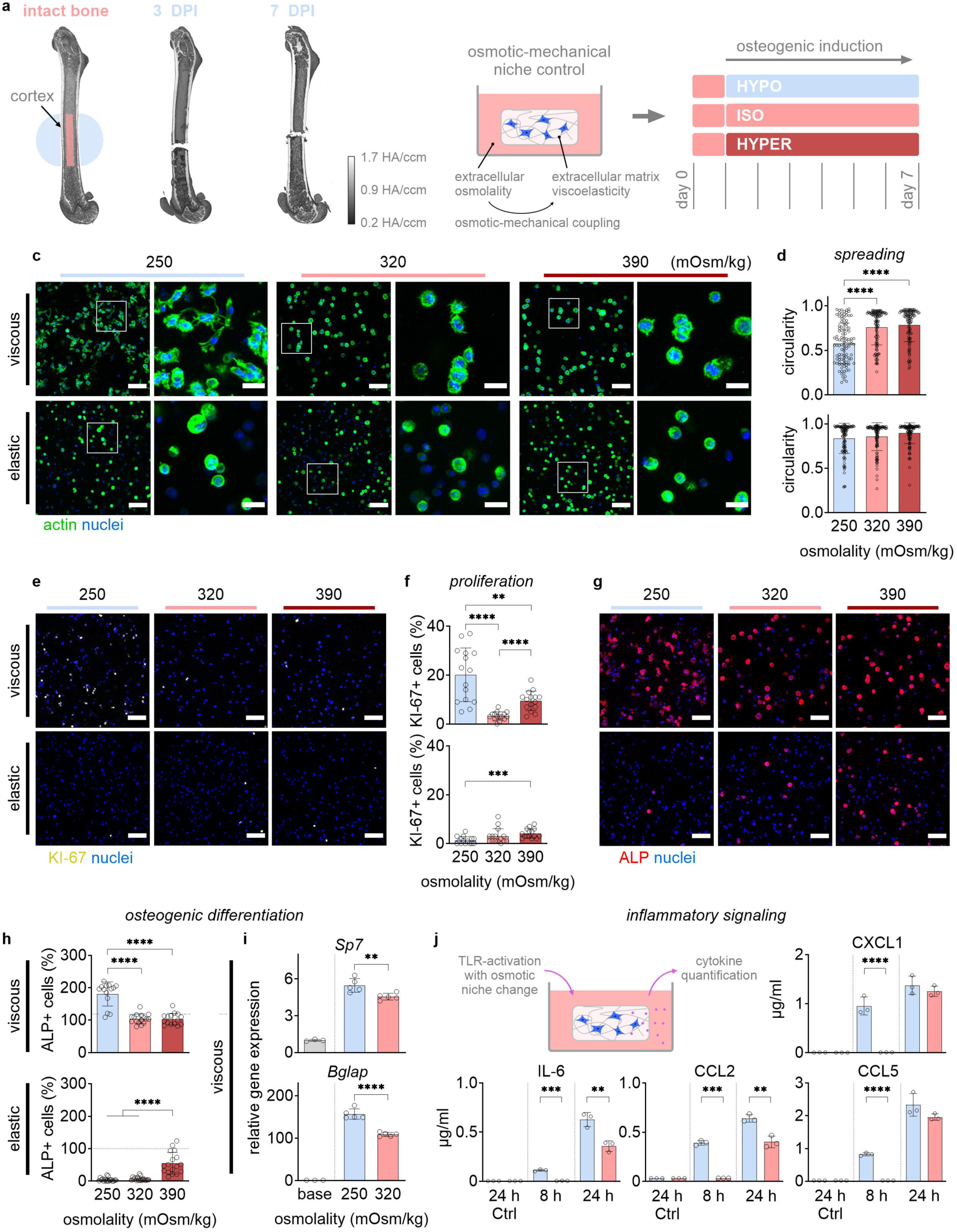
Hypo-osmolality in viscous ECM enhances spreading, proliferation, osteogenic differentiation and cytokine secretion by MSCs. Cell behavior was characterized in terms of spreading, proliferation, and osteogenic differentiation after six days of culture in HYPO (250 mOsm/kg), ISO (320 mOsm/kg), or HYPER (390 mOsm/kg) environments in hydrogels with viscous or elastic mechanical properties. (**a**) Representative uCT images of a mouse bone fracture healing model suggests that the marrow niche would experience an extended period of lowered osmolality after fracture. (**b**) Cell culture protocol used for osteogenic induction of MSCs with modulation of environmental osmolality: osmotic modulation and osteogenic induction started after equilibration to allow cells to attach to adhesion ligands within hydrogels following encapsulation. Cells were cultured in different hydrogel systems with osteogenic induction for up to seven days. (**c**) Representative overview and zoom-in images of nuclei (blue) and actin (green) staining and (**d**) quantification of cell circularity (top: viscous; bottom: elastic) (*n* = 105 single cells from three samples per condition, \*\*\*\**P* < 0.0001 by one-way ANOVA). (**e**) Representative images of nuclei (blue) and KI-67 (yellow) staining and (**f**) quantification of KI-67+ cells (top: viscous; bottom: elastic) (*n* = 15 images from three samples per condition, \*\*\*\**P* < 0.0001, \*\*\**P* < 0.001, and \*\**P* < 0.01 by one-way ANOVA). (**g**) Representative images of nuclei (blue) and ALP (red) staining after 6 days of osteogenic induction and (**h**) quantification of ALP+ cells in a region of interest normalized by the same in ISO condition in viscous ECM (top: viscous; bottom: elastic) (*n* = 14–16 images from three samples per condition in viscous gels, *n* = 17 images from three samples per condition in elastic gels, \*\*\*\**P* < 0.0001 by one-way ANOVA). (**i**) qPCR analysis of osteogenic gene expression under HYPO and ISO condition in viscous ECM at day four of differentiation. The base condition comprised cells equilibrated in viscous ECM, analyzed before osteogenic induction (*n* = 5 samples per condition, \*\*\*\**P* < 0.0001, and \*\**P* < 0.01 by Student’s t test). (**j**) Immune activation assay to study the impact of the osmolality drop on cytokine (purple dots in illustration) secretion by MSCs in viscous ECM under TLR-activation by LPS. Cytokine concentrations were quantified from supernatant collected after 8 h and 24 h. Ctrl conditions were without LPS, HYPO and ISO after 24 h (*n* = 3 samples per condition, \*\*\*\**P* < 0.0001, \*\*\**P* < 0.001, and \*\**P* < 0.01 by Student’s t test). Scale bars: overview 100 µm, zoom-in 30 µm; bar plots: mean ± SD.

The two hydrogel systems described above (Fig. 1f–h, Supplementary Fig. 4) were used as artificial ECM with viscous (fast stress relaxation, *τ*_1⁄2_ = 112 s in ISO) and elastic (slow stress relaxation, *τ*_1⁄2_ > 1000s in ISO) material characteristics^14^. *E* of the viscous (∼14 kPa in ISO) and the elastic (∼23 kPa in ISO) hydrogels were in the respective ranges previously reported to favor osteogenic lineage commitment of MSCs in similar materials^14,34^. In both hydrogel systems, RGD peptides were coupled to alginate chains as ligands enabling cell adhesion to the artificial ECM material. First, cell spreading was quantified by measuring circularity from actin staining (Fig. 2c–d). In viscous hydrogels, circularity was decreased in HYPO compared to ISO, with visibly larger protrusions extending from cells, but unchanged in HYPER conditions. In contrast in elastic hydrogels, circularity was not altered by osmotic changes, which was expected as non-degrading elastic hydrogels restrict cell expansion^34^. Cells could adhere to the elastic, covalently crosslinked hydrogels; this was verified separately through a 2D culture experiment (Supplementary Fig. 6). Proliferation was then quantified by measuring the percentage of KI-67+ cells from staining (Fig. 2e–f). In viscous, proliferation was strongly increased in HYPO, and moderately increased in HYPER, compared to ISO, respectively. In elastic, no difference was observed relative to ISO. The increased proliferation in HYPER in viscous hydrogels suggested a distinct effect of osmolality increased by ion concentration compared to the previously reported decrease in proliferation by applied osmotic pressure in similar materials^35^. The strong increase in cell spreading in HYPO vs. ISO in viscous hydrogels, while changes in hydrogel mesh size could be excluded (Supplementary Note 1, Supplementary Fig. 7), suggested increased cell-matrix interactions, such as ECM remodeling. Increased ECM remodeling of encapsulated cells was previously shown to be relevant in MSCs fate decision^14,36^. The modelled osmolality drop enhanced spreading and proliferation of MSCs in viscous hydrogels with more physiological viscoelastic ECM characteristics.

### ECM stress relaxation directs how osmotic changes regulate lineage commitment

The osteogenic differentiation of precursor cells, such as MSCs, is an essential process in bone regeneration, with alkaline phosphatase (ALP) serving as a key marker of early bone matrix formation and osteoblast differentiation^25,37^. Thus, osteogenic differentiation performance was quantified from ALP staining by ALP+ cells normalized by the ISO condition in (more physiological) viscous ECM (Fig. 2g–h). In viscous ECM, osteogenic differentiation was increased in HYPO compared to ISO. In contrast in elastic, osteogenic differentiation was increased in HYPER compared to ISO. The overall lower differentiation performance in elastic compared to viscous was in line with previous studies that showed faster ECM stress relaxation to increase osteogenic differentiation^14^. Strikingly, our data show that the mechanical characteristics of the ECM (i.e., viscous versus elastic) direct how changes of environmental osmolality regulate early osteogenic differentiation of MSCs. Expression of genes related to osteogenesis was quantified using qPCR after four days of osteogenic induction in viscous ECM. In HYPO compared to ISO, gene expression of essential osteogenic genes was upregulated, including the *Sp7*, encoding for the key osteogenic transcription factor Osterix (Osx), as well as *Bglap*, encoding for osteocalcin in bone matrix formation (Fig. 2i, Supplementary Fig. 8). These results corroborate that the HYPO condition enhanced MSC osteogenic differentiation performance.

### Hypo-osmolality enhances inflammatory signaling

Early pro-inflammatory signaling is essential to initiate the immune-cell recruitment and the angiogenic response that drive successful tissue repair^25^. Inflammatory cues would be expected present in a fresh wound site, such as directly after bone fracture, initiating the pro-inflammatory healing phase. MSCs can respond to inflammatory cues, such as Toll-like receptor (TLR)-activating lipopolysaccharide (LPS)^38^. To assess how the osmolality drop may influence early inflammatory signaling, we quantified cytokine secretion of MSCs stimulated by LPS in different osmotic conditions, while encapsulated in viscous ECM. With LPS, MSCs in HYPO secreted higher levels of CXCL1, CCL2, CCL5, and IL-6 after 8 h, and higher levels of CCL2 and IL-6 also after 24 h, compared to ISO condition (Fig. 2j). In contrast, cytokine secretion without LPS was negligible. The observation that altered incubation osmolality can alter cytokine secretion is in agreement with previous research^39,40^. Our data indicate that a sudden decrease in extracellular osmolality amplifies the secretion of pro-inflammatory and pro-angiogenic cytokines in the presence of an inflammatory cue. Together, these data suggest that an acute osmolality drop upon bone fracture may enhance the early, inflammatory phase in bone healing, potentially facilitating the responses to injury by multiple cell types at the beginning of tissue regeneration. In the following experiments, we focused on MSC osteogenic differentiation processes to unravel how the hypo-osmotic niche change functions as regenerative stimulus.

### Osmolality drop exceeds stress-relaxation effects in proliferation and osteogenic differentiation

In viscous hydrogels, viscoelastic properties were enhanced in decreased osmolality (Fig. 1g). Previous studies reported that faster ECM stress relaxation enhanced cell spreading, proliferation, and osteogenic differentiation of MSCs^14^. To understand the role of altered ECM viscoelasticity in the observed stimulus effect of the osmolality drop on osteogenic differentiation, we focused on comparing the osmotic conditions HYPO and ISO in viscous hydrogels, as viscoelastic matrix properties more closely resemble physiological ECM mechanics. To detect potential changes in mesh size that could influence cell behavior, the hydrogels were analyzed using small-angle x-ray scattering (SAXS). However, no relevant changes in mesh size were observed between HYPO and ISO (Supplementary Note 1, Supplementary Fig. 7); thus, potential influences on cell behavior were excluded. Consequently, the experiments addressed the following: i) the role of cell adhesion to the ECM (i.e., hydrogel), ii) the impact of the osmolality drop on cell behavior through modulating ECM mechanics, and iii) the impact of the osmolality drop on the behavior of encapsulated cells directly (Fig. 3a).

**Fig. 3.**
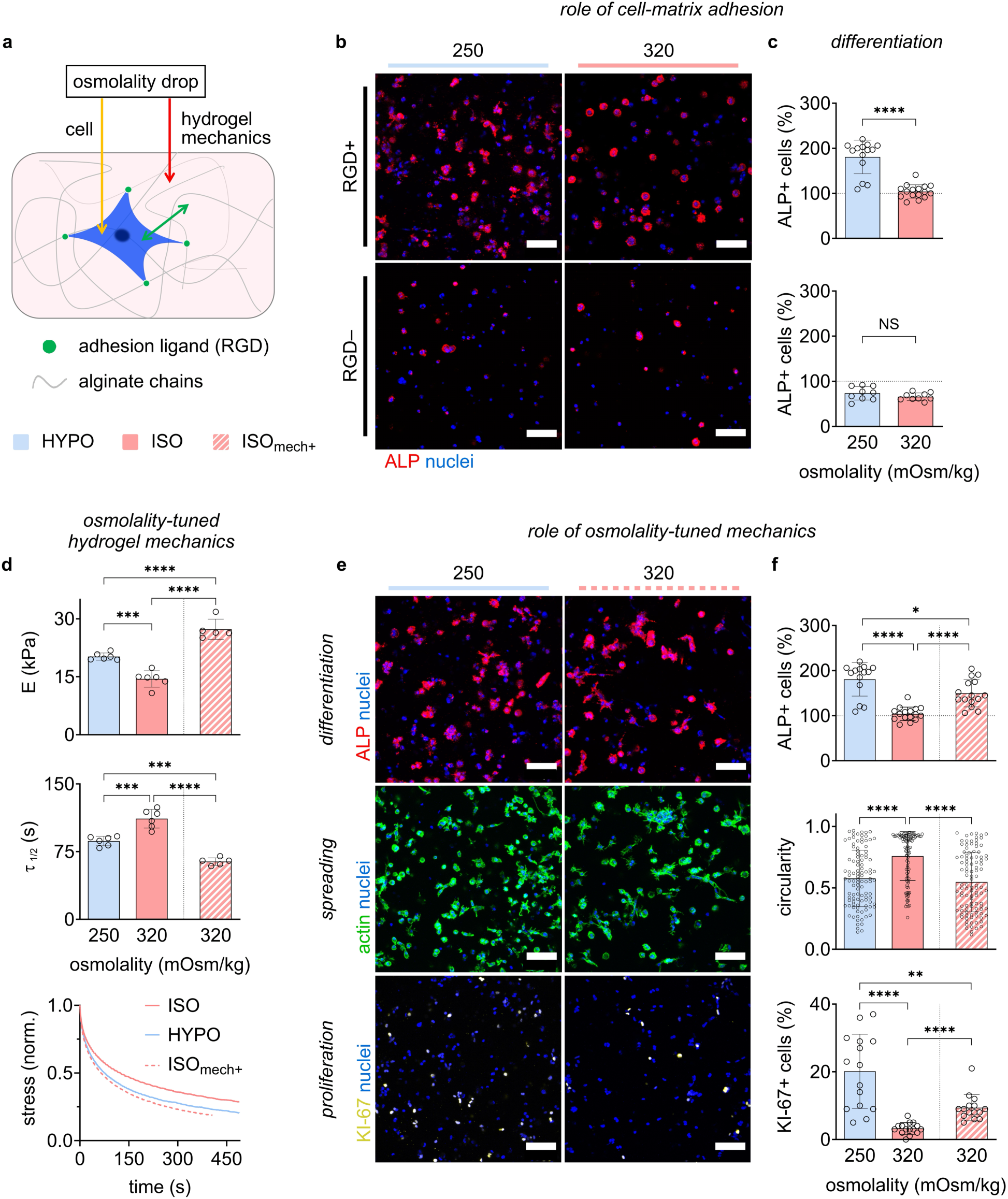
Upregulation of MSC osteogenic differentiation and proliferation by osmolality drop is adhesion-mediated and exceeds effect of altered ECM mechanics. **(a)** Illustration of how the osmolality drop (i.e., lower extracellular osmolality affects ECM mechanics (red arrow) and encapsulated cells directly (yellow arrow). Cell-matrix interactions (green arrow) are mediated by cell adhesion to RGD ligands of the ECM (green dots). Viscous hydrogels with (RGD+) and without (RGD–) RGD adhesion ligands were used to investigate the role of cell adhesion in the stimulating effect of HYPO versus ISO environments. (b) Representative images of nuclei (blue) and ALP (red) staining after 6 days of osteogenic induction with osmotic modulation. (**c**) Quantification of ALP+ cells normalized by the reference (ISO) condition (top: RGD+ gels; bottom: RGD– gels) (*n* = 14–16 images from three samples per condition with RGD+ gels, *n* = 9 images from three samples per condition with RGD– gels, \*\*\*\**P* < 0.0001, and NS *P* > 0.05 by Student’s t test). To isolate effects of the enhanced viscoelastic properties in HYPO compared to ISO condition, hydrogels with increased viscoelastic properties (by altered crosslinking) at ISO osmotic condition were used (ISO_mech+_). (**d**) Mechanical characterization of hydrogels by elastic modulus, *E*, (top), stress relaxation half time, *τ*_1⁄2_, (middle) and representative stress relaxation curves (bottom) (*n* = 5–6 samples, \*\*\*\**P* < 0.0001, and \*\*\**P* < 0.001 by one-way ANOVA). Effects of altered osmotic and mechanical environmental conditions on osteogenic differentiation, spreading, and proliferation were characterized after six days of osteogenic induction. Representative images of staining: (**e**) top: nuclei (blue) and ALP (red); middle: nuclei (blue) and actin (green); bottom: nuclei (blue) and KI-67 (yellow). Quantification of staining: (**f**) top: ALP+ cells in a region of interested normalized by the same in the reference (i.e., ISO) condition (*n* = 14–16 images from three samples per condition, \*\*\*\**P* < 0.0001, and \**P* < 0.05 by one-way ANOVA); middle: cell circularity (*n* = 105 single cells from three samples per condition, \*\*\*\**P* < 0.0001 by one-way ANOVA); bottom: KI-67+ cells (*n* = 15 images from three samples per condition, \*\*\*\**P* < 0.0001, and \*\**P* < 0.01 by one-way ANOVA). Scale bars: 100 µm; bar plots: mean ± SD.

Osteogenic differentiation performance was compared between viscous gels with RGD ligands (RGD+) and without (RGD–) by ALP staining (Fig. 3b). The above-shown increased osteogenic differentiation in HYPO compared to ISO in RGD+, was absent in RGD– (Fig. 3c). This suggests that the stimulus effect of the osmolality drop required cell adhesion to the hydrogels. To quantify effects of the enhanced ECM viscoelasticity in HYPO compared to ISO, we produced control hydrogels with increased viscoelasticity at ISO osmolality (ISO_mech+_), achieved by adapted ionic crosslinking of hydrogels, and compared with the above-used viscoelastic hydrogels (viscous) in HYPO and ISO (Fig. 3d). MSC osteogenic differentiation, spreading, and proliferation were quantified as described above (Fig. 3e). All three parameters were increased in ISO_mech+_ compared to ISO (Fig. 3f), which was in line with previously reported results for faster stress relaxation and higher elastic modulus of the ECM^14^. Strikingly, HYPO showed even higher differentiation performance and proliferation than ISO_mech+_, while cell spreading was similar. This result was even more notable as the ISO_mech+_ hydrogels had faster *τ*_1⁄2_ and higher *E* than viscous hydrogels in HYPO. The osmolality drop increased MSC spreading, proliferation, and osteogenic differentiation with similar characteristics as faster ECM stress relaxation. Together, our results show that the osmolality drop constitutes an adhesion-mediated stimulus for MSC osteogenic differentiation that exceeds the effects of the altered ECM mechanics. The data suggest that the osmolality drop enhances regenerative cell behavior, extracellularly by tuning the mechanics of the ECM, which however, may not account for the entire effect size observed in osteogenic differentiation performance.

### Hypo-osmolality elicits chromatin-associated transcriptional responses in early differentiation

To elucidate the underlying intracellular mechanism, we performed RNA-seq analysis to examine alterations in gene expression linked to our specific cell culture conditions. Given the likelihood of a rapid cellular response to a sudden decrease in extracellular osmolality, as coincides with the biological quest to initiate tissue repair processes, we specifically investigated the early stages of the differentiation process. Therefore, we analyzed different time points following the initiation of osteogenic induction (2h, 12h, 144h) and compared them to the baseline just prior to induction (0h). All experiments were conducted in viscous hydrogels. As baseline, we analyzed the differentiation characteristics over time in the ISO condition (Fig. 4a). Differentially expressed genes (DEGs) showed a continuous progress in gene expression characteristics, as would be expected in the course of cellular differentiation. Clustering by k-means showed four clusters of strongly/mildly up/down regulated DEGs and the respective top five enriched GO terms. The Strong UP cluster was associated with musculoskeletal regeneration (e.g., ossification) and extracellular microenvironment remodeling (e.g., ECM organization), which was in line with histological data reported here, and previously^14^. The Strong DOWN cluster also showed enrichment of positive regulation of cell adhesion, which would appear counterintuitive to the adhesion-mediated mechanosensation in viscous ECM. However, the DEGs leading to enrichment of this GO term were predominantly related to cell-cell adhesion, which would be expected little due to the relatively low cell seeding density in the hydrogels.

**Fig. 4.**
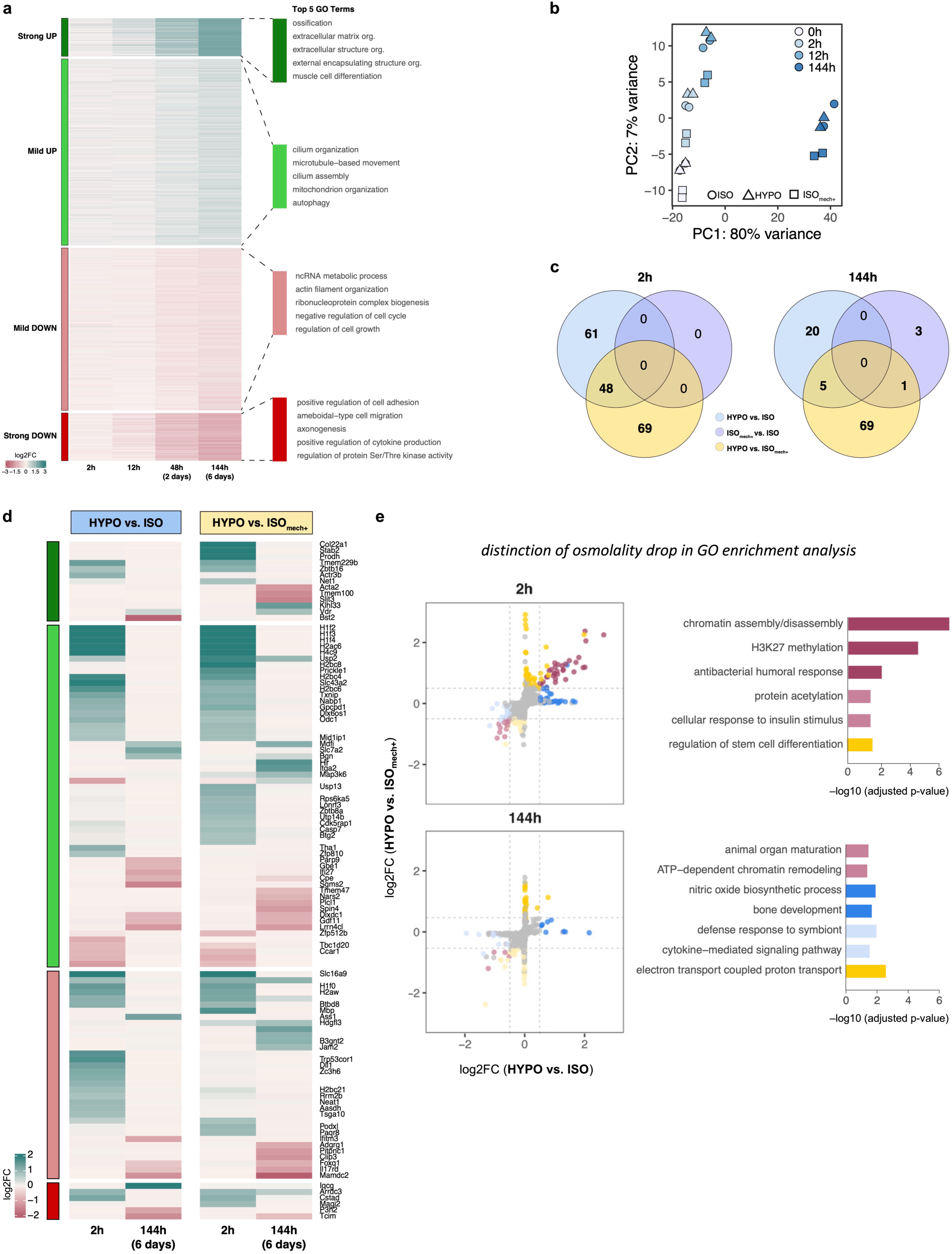
Whole transcriptome sequencing reveals distinct gene expression profile early after osmolality drop. Osteogenic induction of MSCs encapsulated in viscous hydrogels for six days was investigated using RNA sequencing. (**a**) Differentiation of MSCs encapsulated in viscous hydrogels at ISO condition as reference. Differentially expressed genes (DEGs) were analyzed at 2h, 12h, 48h, and 144h (from left to right) after beginning of induction in comparison to directly before (0h). Each row represents a DEG at least one time point. Values are plotted as log_2_ fold change (Log2FC) and clustered by k-means in four clusters (i.e., strong/mild up/down). For each cluster, the top five enriched GO terms are plotted next to the heatmap. (**b**) Principal component analysis (PCA) showing gene expression changes of top 250 regulated genes at the same time points for the control condition 320 (ISO), the osmolality drop condition (HYPO), and the condition with increased viscoelastic properties at ISO osmolality (ISO_mech+_). To better understand differences between gene expression characteristics, we then focused on the early (2h) and late (144h) time points in the differentiation process. (**c**) Global differences between the three conditions (ISO, HYPO, ISO_mech+_) shown as Venn diagram. Each circle represents a pair-wise comparison, and the number of DEGs (either unique or shared) at 2h (left) and 144h (right). Next, the gene expression characteristics in HYPO and ISO_mech+_ conditions, which both enhance cell spreading, proliferation, and osteogenic differentiation compared to the ISO condition, were investigated further. (**d**) DEGs in the comparisons of HYPO or ISO_mech+_ with ISO condition at 2h (left) and 144h (right). DEGs were analyzed based on the cluster information of the baseline differentiation presented in above (A) and plotted similarly as heatmap with Log2FC. Names of top regulated genes are noted on the side of the clusters. Subsequently, we looked outside the baseline clusters and examined the global HYPO-specific changes relative to ISO and/or ISO_mech+_ condition. (**e**) Each dot represents a gene, where all non-grey dots are DEGs in a comparison. Blue dots: DEGs of HYPO versus ISO; yellow dots: DEGs of HYPO versus ISO_mech+_; purple dots: DEGs of HYPO versus both ISO and ISO_mech+_. Darker shades: upregulated genes; lighter shades: down-regulated genes. The top GO terms for each DEG-set are plotted on the right. Top: 2h; bottom: 144h.

To detect when the osmolality drop (HYPO) may have the strongest impact on gene expression compared to ISO, principal component analysis (PCA) was performed for HYPO, ISO, and ISO_mech+_ at the different time points (Fig. 4b). Between HYPO and ISO, PCA showed the most pronounced differences at 2h. Notably, the PCA showed for ISO_mech+_ clear differences to HYPO and ISO at all time points. This was likely caused by the different mechanics of the separate ISO_mech+_ hydrogel batch and the one-day-long equilibration of hydrogels before start of induction. To better understand how effects of HYPO distinguished from ISO, we subsequently focused on comparing 2h versus 144h. To analyze differences in gene expression profiles, we performed pair-wise comparisons which were plotted as Venn diagrams showing the number of DEGs (direct or mutual) between conditions (Fig. 4c). Strikingly, at 2h, HYPO showed 61 and 69 DEGs compared to ISO and ISO_mech+_, respectively, with 48 mutual DEGs. At 144h, this distinction pattern in HYPO was still as pronounced compared to ISO_mech+_, and also, to a lesser degree, compared to ISO. In contrast, these comparisons showed no (2h), or barely any (144h), differences between ISO_mech+_ and ISO. Additionally, we analyzed DEGs of HYPO compared to ISO or ISO_mech+_ based on the cluster information identified for the baseline (i.e., differentiation in ISO, Fig. 4a) (Fig. 4d). The data were plotted as heatmaps together with the names of top regulated genes of the same clusters. The analysis showed similar regulation patterns for both comparisons and underlined the presence of a distinct gene expression profile in HYPO. To analyze how this gene expression in HYPO differs from ISO and ISO_mech+_, global changes relative to the ISO or ISO_mech+_ condition were analyzed and DEGs highlighted in colors (Fig. 4e). Enrichment analysis of the DEG sets revealed that the top-enriched GO term in HYPO was at the early 2h time point ‘chromatin assembly/disassembly’, with involvement of the DEGs H1f0, H1f2, H1f3, H1f4, H2aw, H2bc4, H3f3b, and H4c9. Together, the sequencing data showed that HYPO impacted gene expression in the differentiation process early on and with distinct characteristics compared to ISO and ISO_mech+_ conditions. Notably, genes associated with chromatin structural regulation were more strongly expressed at the early 2h time point. This is an intriguing observation as chromatin structural changes, such as heterochromatinization, have been reported to be involved in gene regulation and fate decision^41–43^. Collectively, these transcriptional data suggest that an acute osmolality drop may promote MSC regenerative processes intracellularly through a fast-acting mechanism associated with chromatin organization.

### Osmolality drop leads to immediate alterations in nuclear morphology, mechanics, and structure

Our RNA sequencing data showed that genes related to chromatin structure changes were altered already two hours after osmolality drop (HYPO). Considering that such alterations in gene expression alterations likely are only detectable with some delay, this suggests a fast impact of the osmolality drop on nuclear structures. To understand better how extracellular hypo-osmolality affected cellular behavior in our experiments, we characterized nucleus morphology first. We analyzed cells incubated for just 15 minutes with osteogenic induction stimulus and different osmolality inside hydrogels using high-resolution confocal imaging of nucleus staining. Nucleus volume (Fig. 5a) as well as nucleus sphericity (Fig. 5b) were higher in HYPO compared to ISO condition. These observations are in agreement with previous studies^44^, and suggest that decreased extracellular ion concentration had a rapid impact on nucleus biophysical properties. Thus, we next analyzed mechanical properties of the nucleus. We used Brillouin microscopy to measure maps of the Brillouin shift, a proxy readout of material mechanics related to compressive modulus, in cross sections of nuclei in live cells after 15 to 60 minutes in HYPO and ISO incubation. Curiously, the mean Brillouin shift in nuclei decreased with more hypo-osmolar incubation conditions, indicating softening of nuclear structures in HYPO (Fig. 5c–d). This observation is in agreement with previous research reporting nucleus softening in hypo-osmolality^45^, and interesting also considering that softening of nuclear structures has previously been related to decreased chromatin condensation^46,47^.

**Fig. 5.**
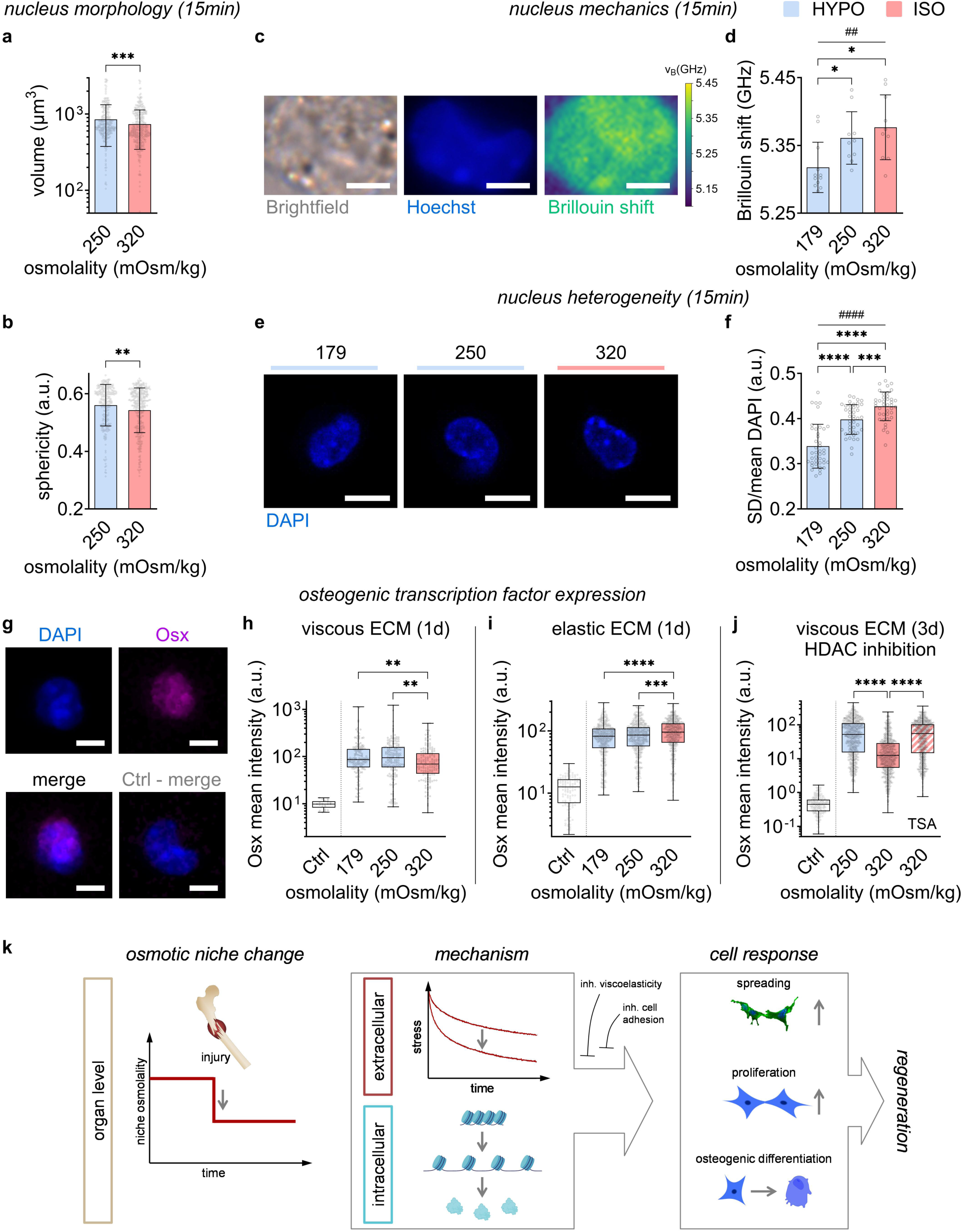
Osmolality drop causes immediate changes in nucleus morphology, mechanics, and heterogeneity, and enhances osteogenic transcription factor expression early on. **(a)** Nucleus volume and (b) nucleus sphericity were quantified for cells in viscous ECM fixed after 15 minutes incubation with osteogenic induction in HYPO or ISO conditions (*n* = 261–356 single cells per condition, \*\*\**P* < 0.001, and \*\**P* < 0.01 by Mann-Whitney test). To assess nucleus mechanical properties in HYPO and ISO incubation, with osteogenic induction, live cells in viscous ECM were imaged after 15 to 60 minutes incubation using Brillouin microscopy. (**c**) Representative images of an imaged cell nucleus from brightfield microscopy, fluorescence imaging of Hoechst staining, and a map of Brillouin shift (scale bars: 4 µm). (**d**) Quantification of the Brillouin shift, a readout relating to material elastic modulus, mean value across a nucleus, indicating nucleus softening quickly after the osmotic change (*n* = 10–12 single cells per condition, \**P* < 0.05 by one-way ANOVA, ##*P* < 0.01 by Spearman’s rank correlation). To analyze a potential impact of HYPO on chromatin condensation, we analyzed the heterogeneity of nuclear staining, DAPI, on cells fixed after 15 minutes of incubation in osteogenic induction with HYPO and ISO in viscous ECM. (**e**) Representative images of DAPI staining in 179, 250, and 320 mOsm/kg conditions (scale bars: 10 µm). (**f**) Heterogeneity of nuclei was quantified by the ratio between standard deviation (SD) and mean value of DAPI signal (*n* = 39–43 single cells from three samples per condition, \*\*\*\**P* < 0.0001, and \*\*\**P* < 0.001 by one-way ANOVA, ####*P* < 0.0001 by Spearman’s rank correlation). Transcription factor (TF) expression was analyzed after culture with osteogenic induction in HYPO and ISO conditions to elucidate the impact of the acutely lowered extracellular osmolality on osteogenic differentiation by MSCs, focusing on Osterix (Osx) staining in the nucleus. (**g**) Representative images of DAPI and Osx staining (top), and merged images (bottom, left); a merged image of a negative control (secondary antibody only) shows no detectable Osx signal (bottom, right). Osx expression was quantified as mean nuclear signal intensity. (**h**) Quantification after one day shows increased Osx expression in HYPO conditions compared to ISO in viscous ECM (*n* = 154–178 single cells per condition, \*\**P* < 0.01 by Kruskal-Wallis test). To study the role of ECM viscoelasticity in the observed effects, we performed the same experiment using elastic instead of viscous ECM. (**i**) Contrasting observations in viscous ECM, Osx expression was decreased in HYPO compared to ISO in elastic ECM (*n* = 509–615 single cells per condition, \*\*\*\**P* < 0.0001, and \*\*\**P* < 0.001 by Kruskal-Wallis test). To test whether changes in chromatin organization under HYPO in viscous ECM, which were suggested by analyses of transcriptome sequencing, nuclear mechanics and nuclear structures, could have caused the increase in Osx expression, we supplemented ISO medium with histone deacetylase (HDAC) inhibitor trichostatin A (TSA) at 75 nM concentration to cause chromatin decompaction. (**j**) After three days of osteogenic induction in viscous ECM, Osx expression with TSA was higher than in standard ISO medium (both 320 mOsm/kg), and at similar levels as in HYPO condition (250 mOsm/kg) (*n* = 750–839 single cells per condition, \*\*\*\**P* < 0.0001 by Kruskal-Wallis test). Bar plots: mean ± SD. (**k**) Schematic representation of the proposed mechanisms underlying the osmolality drop as bone regenerative stimulus upon fracture. At organ level, marrow osmolality is suddenly decreased due to the fracture of the cortical bone barrier. The osmotic niche change increases bone regenerative cellular processes: extracellularly by enhancing matrix stress relaxation, and intracellularly, by decreasing chromatin compaction and enhancing transcription factor expression. This leads to altered gene expression and protein synthesis, steering the cellular response to injury by niche resident multipotent cells. The regenerative effect of the osmotic niche change depends on the viscoelastic and adhesion properties of the ECM; inhibition of either factor abolishes the regenerative effect.

Chromatin condensation (i.e., compaction) is regulated reversibly by ions interacting with the charged chromatin structures^48,49^. A change in extracellular ion concentrations would be expected to affect ion concentration gradients between intracellular and extracellular space. Such alterations could cause ion efflux from the cell, and consequently from the nucleus as nuclear pore complexes are permeable to ions^50^. To elucidate whether hypo-osmolality in our 3D culture system might induce chromatin decondensation, we quantified signal heterogeneity of nuclear staining 15 minutes after the osmolality drop in viscous hydrogels to detect potential changes in nuclear structures. Heterogeneity of DAPI staining decreased with decreasing incubation osmolality (Fig. 5e–f, Supplementary Fig. 9), already after 15 minutes, which suggested that the osmolality drop decreased chromatin condensation. This was in line with previous reports of chromatin decondensation in hypo-osmolality^44,51^.

### Hypo-osmolality enhances osteogenic transcription factor expression

Our data show that hypo-osmolality induced changes in nucleus biophysical properties in agreement with previous publications on nucleus morphology and mechanics under osmotic changes^44,52^, charge-mediated intermolecular interactions between macromolecules^53,54^, such as DNA^48,49^, and chromatin condensation^55^. Considering that chromatin compaction is known to affect gene regulation, and assuming that acutely lowered osmolality in HYPO decreased chromatin compaction, we hypothesized that a potential related impact on MSC differentiation processes would occur early on. Thus, we quantified staining of the transcription factor (TF) Osterix (Osx) in the nucleus in different conditions with osteogenic induction (Fig. 5g). Osx is an essential TF in differentiation of mesenchymal progenitors into osteoblasts. Strikingly, Osx expression after one day was higher in hypo-osmolar (179 and 250 mOsm/kg) compared to ISO conditions (320 mOsm/kg) in viscous ECM (Fig. 5h). This corroborates that the osmolality drop has a fast impact on MSC osteogenic differentiation processes. To test the role of physiological ECM viscoelasticity properties in the impact of the osmolality drop, we perturbed the experiment using non-physiological elastic ECM. Contrasting observations of cells in viscous ECM, Osx expression after one day was lower in hypo-osmolar compared to ISO conditions in elastic ECM (Fig. 5i). Together, these data suggest that ECM stress relaxation properties directed the cellular response to the same osmotic change, HYPO. This observation appears in line with the above reported distinct effects of HYPO or HYPER conditions on osteogenic differentiation performance after six days culture in viscous or elastic ECM (Fig. 2g–h).

Considering that HYPO enhanced Osx early on, we asked whether indeed chromatin decompaction could have caused the higher Osx expression. Previous studies reported that extracellular hypo-osmolality can influence cell differentiation, decreasing chromatin condensation and altering transcription^51^. Therefore, we used the small molecule drug trichostatin A (TSA), which inhibits histone deacetylases (HDACs) and decompacts chromatin^56,57^. We used the same quantification approach of Osx staining as described above to compare HYPO and ISO conditions to ISO supplemented with 75 nM TSA after three days of osteogenic induction (Fig. 5j). Again, HYPO showed higher Osx staining than ISO, in agreement with day one. Remarkably, also the iso-osmolar TSA condition showed higher Osx compared to ISO, at similar levels as HYPO. This observation is in agreement with a previous study reporting that TSA enhanced MSC osteogenic differentiation, showing increased expression of the osteogenic TF Runt-related transcription factor 2 (Runx2)^58^, another key TF in osteoblast differentiation. We thus also quantified Runx2, and as well observed higher Runx2 expression in both HYPO and iso-osmolar TSA-supplemented conditions compared to ISO (Supplementary Fig. 10). These results are in agreement with a previous study showing that extracellular hypo-osmolality can influence cell differentiation by chromatin decondensation^51^. Together our analyses of nucleus biophysical properties and TF expression suggest that the sudden hypo-osmolar niche change in marrow upon bone fracture functions in a fast-acting manner through decompacting chromatin, enhancing osteogenic TF expression to promote bone regenerative processes by MSCs.

## Discussion

This work uncovers osmotic niche disruption as a fast-acting physical signal that links tissue injury to regenerative cell programs. Ex vivo analyses of human tissue suggest that fracture of the cortical bone boundary exposes the marrow niche to a rapid drop in osmolality. This osmotic shift enhances ECM stress relaxation in fracture haematoma, blood clots, and osmotically tunable hydrogels (Fig. 1), and promotes MSC spreading, proliferation, osteogenic differentiation, and inflammatory cytokine secretion in a 3D viscoelastic matrix (Fig. 2). The regenerative response requires cell–ECM adhesion and matrix stress relaxation, indicating that ECM mechanics gate cellular interpretation of osmotic cues (Fig. 2, Fig. 3). Rapidly, the osmolality drop alters nuclear morphology, mechanics, and chromatin organization, preceding increased osteogenic transcription factor expression (Fig. 4, Fig. 5). These findings suggest that nuclear structures rapidly respond to disruptions in extracellular osmotic–mechanical cues, providing a physical pathway by which injured niches coordinate ECM mechanics and nuclear state to initiate regenerative programs (Fig. 5k).

Osmotic control of ECM viscoelasticity also reframes how artificial matrices should be designed. Rather than treating ion concentration, charge density, and water binding as background formulation variables, our data suggest that these parameters can be used to program stress relaxation and cellular responses. This principle extends previous observations in ex vivo tendon^10^ to fracture-associated matrices and synthetic hydrogels (Fig. 1), indicating that osmotic–mechanical coupling may be a transferable feature of charged extracellular materials. Designing biomaterials around this coupling could improve 3D models of regeneration and disease and provide new ways to tune cell–matrix interactions.

A second implication is that osmotic niche disruption is rapidly coupled to nuclear structure in 3D viscoelastic matrices. The osmolality drop altered nuclear morphology, mechanics, and DAPI heterogeneity within minutes, while RNA sequencing revealed early enrichment of chromatin-associated programs, including linker histone H1 genes. Because H1 regulates chromatin structure^59^ and is sensitive to ionic conditions^60^, as is chromatin compaction, these findings suggest that extracellular osmotic changes may rapidly alter nuclear organization and transcription factor access. Notably, H1 has also been implicated in T cell activation, suggesting relevance beyond MSC differentiation^61^. The dependence of the HYPO stimulus on ECM stress relaxation and cell–ECM adhesion may reflect that ECM mechanics shape cytoskeletal organization, which influences membrane tension and ion-channel-mediated volume adaptation during osmotic challenge^11,62^. Further study will be required to define how osmotic niche disruption is transmitted intracellularly to regulate nuclear state and transcriptional activation.

Because many pathologies involve dysregulation in matrix mechanics, charge properties, or fluid content, osmotic–mechanical niche changes may extend beyond bone fracture. Inflammation and edema alter solute and water exchange^4,63,64^, fibrosis and cancer remodel ECM stiffness and viscoelasticity^28,65,66^, and changes in glycosaminoglycan sulfation may alter tissue charge properties in osteoarthritis, Alzheimer’s disease, and cancer^67^. Our findings suggest that such pathological microenvironments should be studied not only through stiffness or biochemical signaling, but also through integrated osmotic, charge, and viscoelastic cues in 3D matrices. Incipient coagulation may further reshape osmotic–mechanical cues after injury by altering local ion binding and extracellular charge density^68–71^.

Our finding that hypo-osmolality enhances MSC osteogenic differentiation is broadly consistent with prior work linking diluted medium to increased osteogenesis^35^. However, our injury-inspired protocol applied hypo-osmolality at induction onset, accounted for ECM mechanics, and showed that PEG-based osmotic compensation did not restore ISO-like differentiation, indicating a mechanistically distinct regenerative stimulus (Supplementary Note 2, Supplementary Fig. 11).

Together, these findings establish osmotic niche disruption as a fast-acting physical mechanism by which injury converts compartment-level osmotic differences into coordinated ECM and nuclear responses that initiate regenerative programs. By linking niche osmolality changes to matrix viscoelasticity, chromatin organization, and cell fate, this work expands the cellular niche concept beyond biochemical and mechanical cues to include osmotic–mechanical state changes as regulators of regeneration. More broadly, it suggests that tissue boundary disruption can generate immediate physical signals that shape coordinated multicellular responses after injury. Future studies should determine whether similar osmotic–mechanical mechanisms operate across regenerative, inflammatory, and pathological tissue remodeling contexts, and how they can be leveraged in novel therapeutic approaches.

## Methods

Human tissue sample harvesting was approved by the Ethics Board of the Charité - Universitätsmedizin Berlin (EA1/125/10, EA1/194/13). All subjects provided written informed consent. Human fracture haematoma samples were obtained from five patients (four female, one male; median age of 66 years) undergoing surgery procedures that required the removal of the tissue. Samples of bone marrow, peripheral blood, and cerebrospinal fluid as well as of fracture haematoma were collected from adult donors without any conditions that should influence sample quality.

All animal procedures complied with the German Animal Welfare Act and EU Directive 2010/63/EU and were approved by the State Office of Health and Social Affairs Berlin (LAGeSo), approval no. G0260/18. To keep the number of animals to a minimum, the samples were taken from other research projects but with no relationship that would affect tissue properties in the context of this study.

### Ex vivo tissue osmolality measurement

The osmolality of a sample was measured using a freezing-point osmometer (OSMOMAT Auto, Gonotec). Ex vivo samples of peripheral blood serum, bone marrow plasma, and cerebrospinal fluid were measured. All samples were stored at -80 °C after harvesting before thawing for subsequent osmolality measurement at standard room temperature.

### Blood clot casting

Artificial blood clots were casted using peripheral blood of a healthy individual (female, 26 y) and in cylindrical molds of eight mm diameter, similar to casted hydrogels. Before casting, blood was stored in citrate tubes. To achieve homogenously coagulated samples in the molds, coagulation during the mixing and casting steps was avoided by pre-cooling molds, pipette tips and blood. Coagulation was achieved by mixing blood with a solution of thrombin and CaCl_2_ (Tisseel, Baxxter) at a concentration of 9.1% of the final sample volume. After casting, molds were placed in a petri dish covered by a lid for ten min in an incubator to achieve full coagulation of the samples.

### Osmotic environment modulation

To investigate the influence of altered osmotic environments on biophysical properties of ex vivo tissue samples and hydrogels, different osmotic environments were established by altering the concentration of the buffer surrounding a sample. Ex vivo tissue samples, after harvesting and dissection, and artificial blood clots, were incubated in different osmotic buffers for five to ten hours. A fracture haematoma of a donor was divided into comparably sized samples, which were then incubated in different osmotic conditions. Hydrogels first were equilibrated in iso-osmolar buffer (DMEM) for one day and then incubated in different osmotic buffers overnight.

For fracture haematoma samples and blot clots, ddH_2_O and phosphate buffered saline (PBS) without Ca^2+^ and Mg^2+^ (D8537, Sigma) were used to establish HYPO and ISO environments, respectively. For hydrogels, defined osmotic environments were established using formulations suitable for cell culture, as described below. As buffer for material characterizations of the hydrogels, Dulbecco′s Modified Eagle′s Medium (DMEM) was used as buffer instead of PBS for better comparability to cell culture experiments (cat. # D5546, Sigma). For HYPO conditions, the DMEM concentration was decreased by dilution with ddH_2_O. For HYPER conditions, higher DMEM concentrations were established by diluting 10x concentrated DMEM (cat. # D2429, Sigma) with ddH_2_O. Osmolality of osmotic environments was measured with the method used also for ex vivo tissue samples, as described above. In the hydrogel-based 3D culture system, the ratio of medium (V_medium_) to sample (V_sample_) volume was found to potentially affect the mechanical properties of a sample (Supplementary Fig. 4) and thus was kept constant at 8 throughout the study.

### Alginate hydrogel preparation

Alginate hydrogels were produced at a final concentration of 20 mg per ml. Casting was performed between glass slides and cylindrical samples of eight mm diameter were achieved using a biopsy punch. Integrin-binding ligands were established on the alginate chains by covalently coupling RGD peptides at a concentration of 112.2 mg per g alginate. For viscoelastic hydrogels mimicking osmo-mechanical interplay characteristics, RGD peptides (GGGGRGDSP, Peptide 2.0) were coupled to 1 g low molecular weight alginate (cat. # 4200501, Novamatrix) by carbodiimide chemistry using 274 mg N-hydroxysuccinimide (cat. # 130672, Sigma) and 484 mg 1-ethyl-3-(3-dimethylami-nopropyl)-carbodiimide hydrochloride (cat. # E7750, Sigma). The reaction was conducted for 20 h with constant stirring in 0.1M MES hydrate (cat. # M8250, Sigma), 0.3M NaCl buffer at pH 6.5 and stopped by hydroxylamine hydrochloride (cat. # 26103, Thermo Fisher) at 36 mg per ml. Subsequently, dialysis was performed with 3.5 kDa MWCO tubing (cat. # 11495869, Spectrum) for at least 4 days in decreasing NaCl concentrations in ddH_2_O, finishing with 0 g/L over 2 days, with 3–4 changes per day. After dialysis, the alginate solution was purified using activated charcoal (Sigma) and was sterile filtered (cat. # SCGP00525, Merck). Finally, the sterile solution was freeze-dried for storage at -20 °C until use for hydrogel production. For crosslinking of the alginate chains, 1.22 M CaSO_4_ slurry was used similar to the method described by Chaudhuri et al.^14^ and at concentrations of 5% and 7% for standard and ‘mech+’ conditions, respectively. For elastic gels, first, norbornene-functional groups were coupled to high molecular weight alginate (cat. # 4200101, Novamatrix) via carbodiimide chemistry, similar to the method described by Lueckgen et al.^72^. To achieve RGD binding sites on the alginate chains, a thiol-containing sequence (CGGGGRGDSP, Peptide 2.0) was coupled in the same UV-initiated reaction with thiol-ene click chemistry as the crosslinking. For coupling of RGD to alginate chains and the crosslinking between alginate chains, the photoinitiator Irgacure (cat. # H1361, TCI) and the crosslinker DL-Dithiothreitol (cat. # 43816, Sigma) were added at concentrations of 5 mg per ml and 0.3 mg per ml, respectively, to a mix of alginate and RGD peptide in PBS without Ca^2+^ and Mg^2+^ (cat. # D8537, Sigma). Crosslinking between alginate chains and coupling of RGD peptide to alginate was initiated by exposure to 365 nm UV light for 1 min at 20 mW/cm^2^ (OmniCure S2000) in a custom-built chamber.

### Mechanical characterization

Mechanical characterization of fracture haematoma, blood clots and hydrogels was performed using unconfined uniaxial compression testing (TestBench LM1 system, BOSE). For all measurements, a 250 g load cell (Model 31 Low, Honeywell) was employed. Sample diameter was measured using calipers and height using the BOSE system by lowering the top plate until contact. All hydrogel samples were measured as discs of 8 mm diameter and 2 mm height. Fracture haematoma samples and blood clots were prepared to match comparable dimensions. The testing was performed similar to the protocol reported by Chaudhuri et al.^14^. In brief, the compression was carried out between two parallel plates (one fixed, one moved in displacement-controlled mode in z-direction) at 0.016 mm/s and until 15 % strain and, subsequently, held constant to record decreasing loads due to material stress relaxation behavior. A MATLAB script was used to calculate *E* and *τ*_1/2_. *E* was calculated from a 5 % strain interval in the linear region of the generated stress-strain curve. *τ*_1/2_ was defined as the elapsed time at maximum compression until half of the peak stress was released.

### Swelling characterization

Hydrogel swelling was characterized volumetrically and gravimetrically after incubation in different osmotic (i.e., DMEM buffer) concentrations overnight. After producing a ∼2 mm thick gel film, gels had been punched out by an 8 mm diameter biopsy punch and incubated subsequently. Size measurement by diameter (caliper) and thickness (TestBench LM1 system, BOSE) showed no differences between osmotic conditions for the used hydrogels. Wet weight measurements showed only small differences (∼1%) in the extreme comparison of hypo-osmolar ddH2O versus iso-osmolar DMEM (Supplementary Fig. 2).

### Differential scanning calorimetry

Thermoanalytical characterization was performed using DSC with an EXSTAR DSC 7020 (SII Nanotechnology Inc.). T_melt_ was quantified for probes of ex vivo fracture haematoma and viscous hydrogels (with ionic crosslinking), which mimic the viscoelastic properties of fracture haematoma. In the hydrogels, changes in W_f_ due to osmotic modulation were quantified.

### ICP-MS measurements

Calcium ion concentration in hydrogels was quantified using inductively coupled plasma mass spectrometry (ICP-MS). The measurements were performed at the German Federal Institute for Risk Assessment (BfR).

### Mesh size analysis using SAXS

SAXS was used to investigate potential alterations in hydrogel mesh size of ionically crosslinked hydrogels (viscous) after incubation in ddH_2_O, HYPO, or ISO solution. The measurement data provided information on the structure of the gel in the size range 40 nm to 0.5 nm. SAXS measurements were performed in a gel sample holder with a Kratky-type instrument (SAXSess from Anton Paar, Austria) at room temperature. The SAXSess instrument has a low sample-to-detector distance of 0.309 m. This is appropriate to investigate materials with low scattering intensities. Conversion to absolute scale was performed for the measured intensity as described by Orthaber et al.^73^. The scattering vector q is defined in terms of scattering angle *θ* and the wavelength *λ* of the radiation (*λ* = 0.154 nm): thus, *q* = 4π*n*/*λ* sin *θ*. Deconvolution (slit length desmearing) of the SAXS curves was performed with the SAXS-Quant software. Samples analyzed with SAXS were used as prepared. The method is described in further detail in Supplementary Note 1.

### µCT of bone fracture healing

Representative ex vivo micro-computed tomography (µCT) images illustrating a typical course of fracture healing were sourced from an independent murine femoral osteotomy dataset in 10-week-old female C57BL/6J mice. A mid-diaphyseal transverse osteotomy (∼0.68 mm) was created in the left femur and stabilized with a four-pin plate fixator. Only sham-handled animals (identical anesthesia/handling without mechanical stimulation) were used. Femora were scanned unoperated (intact bone) or postoperatively, three (3 DPI) or seven (7 DPI) days post-osteotomy. Scans were acquired ex vivo using a vivaCT 40 system (Scanco Medical; 10.5 µm isotropic voxel size, 55 kVp, 145 µA, 300 ms). Images are shown solely to qualitatively depict the fracture gap and typical callus maturation; no µCT-based quantification or group comparisons are reported in this work.

### Cell culture with osmotic modulation

Cells of the mMSC D1 cell line (cat. # CRL-12424, ATCC) were used for cell culture experiments. Cells were expanded in cell culture flasks and maintained at sub-confluency before encapsulation into the hydrogels. For main experiments, cells were used below passage six. For osmotic modulation, a medium formulation was developed based on a standard expansion medium for MSCs, consisting of DMEM (cat. # D5546, Sigma) with 10% FBS (cat. # S0615, Sigma), 1% Pen/Strep (cat. # A2213, Merck), 1% GlutaMAX (cat. # 35050061, Gibco), with osteogenic induction media containing 50 µM L-ascorbic acid (cat. # A8960, Sigma), 10 mM ß-glycerophosphate (cat. # G9422, Sigma), and 0.1 µM dexamethasone (cat. # D2915, Sigma). The osmolality of media was tuned by adjusting the DMEM concentration in the buffer part of the medium mix, as illustrated in Supplementary Fig. 5a. The pH values of osmolality-modulated cell culture media were at a similar level as measured for standard ISO medium.

### Encapsulation of MSCs in hydrogels

Cells were encapsulated at a concentration of 3×10^6^ cells per ml hydrogel. Hydrogels were casted as described above and at a sample height of 1 mm. Gels were equilibrated after encapsulation in iso-osmolar expansion medium overnight to allow the cells to adhere to RGD adhesion ligands. Next, osteogenic induction medium was applied, with consistent supplement concentrations (Supplementary Fig. 5a), while the osmolality of the culture medium was modulated, as illustrated in Supplementary Fig. 5b.

### Cell viability quantification

Viability of cells encapsulated in hydrogels was quantified after 2 days of culture in a wide range of chosen osmotic conditions (38–461 mOsm/kg). Gels were stained with calcein AM (1:1000) and propidium iodide (1:2500) for 15 min in culture medium. For quantification, at least 3 ROIs per gel were imaged with 50 µm thick z-stacks using live-cell microscopy (Leica). ImageJ was used to apply maximum projection to z-stacks and count live (LC) and dead (DC) cells. Viability was calculated by *viability* (%) = *LC*⁄(*LC* + *DC*).

Metabolic activity was quantified after 2 days culture and normalized by the baseline at day 0. In brief, gels were incubated for 1 h in 1:10 mix of alamarBlue (Bio-Rad) and culture medium. Subsequently, fluorescence was measured using a plate reader (Tecan).

### Immunohistochemistry

Cells inside gels were fixed using 4% paraformaldehyde and permeabilized using 0.1 % Triton-X100 and 0.1 % Tween-20 for 10 min in DMEM. Subsequently, gels were stabilized for further processing steps using 100 mM HEPES buffer with 5 mM BaCl_2_ and stored in PBS at 4 °C until staining. Nuclei were stained with DAPI (cat. # D1306, Invitrogen) or Draq5 (cat. # 424101, BioLegend); actin with Alexa Fluor-488 Phalloidin (cat. # A12379, Invitrogen); proliferating cells with rabbit anti-KI-67 (cat. # ab16667, Abcam) and goat anti-rabbit Alexa Fluor-647 (cat. # A21246, Invitrogen); alkaline phosphatase with ELF97 (cat. # E6588, Invitrogen). TF expression was analyzed using rabbit anti-Sp7/Osterix (cat. # ab209484, Abcam) and mouse anti-Runx2 (cat. # ab76956, Abcam) followed by incubation with actin and nucleus staining along with secondary antibodies, such as goat anti-rabbit Alexa Fluor-647 (cat. # A21246, Invitrogen) and goat anti-mouse Alexa Fluor-555 (cat. # A21422, Invitrogen). Nuclei in live cells were stained using Hoechst (cat. # H3570, Invitrogen). Stainings were applied following standard protocols for immunohistochemistry.

### Confocal microscopy

During imaging, hydrogel samples were kept hydrated to avoid evaporation. Regions of interest (ROIs) were chosen with sufficient distance to the hydrogel surface to avoid distorting effects on cell behavior. The same settings were used for collecting images across all conditions within an experiment. Unless described otherwise, ROIs were imaged using a SP5 II confocal microscope (Leica) with either a 25X or a 63X water immersion objective. ImageJ software was used to quantify imaging data.

### Image analysis

Histological markers analyzed after six days of osteogenic induction, including cell spreading (phalloidin), proliferation (KI-67) and osteogenic differentiation (ALP), were quantified from maximum projections of z-stacks. Cell spreading was quantified by cell circularity. Cell proliferation was quantified by the percentage of KI-67-positive nuclei and the overall number of nuclei in the ROI after 7 days cell culture. Osteogenic differentiation was quantified by the number of ALP-positive cells in the ROI. For comparison of osteogenic differentiation performance, the number of ALP-positive cells was normalized to the reference condition of viscoelastic gels, which mimic the mechanical condition in fracture haematoma, in iso-osmolar condition (ISO).

Nucleus morphology was analyzed 15 minutes after the beginning of the osmotic environment modulation. Three-dimensional nuclear morphology was quantified from confocal z-stacks of DAPI-stained samples. For each segmented nucleus, volume and sphericity were computed. Nuclei touching image borders were excluded. For data analysis, outliers identified by ROUT method were excluded in GraphPad PRISM.

Nucleus heterogeneity, used to detect potential changes in chromatin condensation, was analyzed 15 minutes after the beginning of the osmotic environment modulation. DAPI was used to stain nuclei. ROIs were imaged by confocal z-stacks. Heterogeneity of staining signal inside a nucleus was then analyzed using ImageJ, and quantified by the ratio between standard deviation (SD) and mean value of the signal, as previously reported by Lima et al.^51^.

Transcription factor expression was quantified from confocal z-stacks of stained samples by integrating fluorescence intensity within each nucleus. Nuclear masks generated from DAPI served as ROIs for extracting intensity values from the TF fluorescence channel. For each nucleus, total integrated density and mean fluorescence were calculated. Mean nuclear TF intensity was used for comparisons, as it normalizes TF signal to nuclear area and allows assessment of per-nucleus expression independent of nucleus size.

### Brillouin microscopy

Nucleus mechanical properties were assessed by Brillouin measurements in live cells encapsulated in hydrogels, 15 to 60 minutes after the beginning of the osmotic environment modulation. Measurements were performed using an inverted confocal Brillouin microscope (Discoverer from CellSense, Berlin, Germany). The microscope setup featured an excitation wavelength of 780.24 nm and was equipped with a 40X C-Apochromat 1.20/W water immersion objective (Zeiss). Samples were measured at 37 °C.

### Cytokine secretion assay

MSCs were encapsulated in hydrogels as described above and then cultivated in HYPO or ISO medium supplemented with 0 or 100 ng/ml LPS for TLR-activation. Supernatants were collected at 8 h and 24 h after stimulation, centrifuged at 1000 x g for 10 minutes at 4°C, and stored at −80 °C until analysis. Quantification of cytokine levels was performed using LEGENDplex^TM^ multiplex immunoassays (cat. # 741026 and cat. # 740621, Biolegend) according to the manufacturer’s protocol. A Sony ID7000 flow cytometer was used to analyze the samples. Measurement data were processed using LEGENDplex^TM^ Data Analysis Software Suite (BioLegend).

### qPCR

To compare the expression of genes related to osteogenic differentiation of MSCs between HYPO and ISO conditions, qPCR was performed. After RNA isolation from hydrogels using the RNeasy Micro Kit (cat. # 74004, Qiagen), the RevertAid First Strand cDNA Synthesis Kit (cat. # K1621, Thermo Fisher) was used for reverse transcription. qPCR was performed using PowerTrack SYBR Green PCR Master Mix (cat. # A46113, Applied Biosystems) and the Quantstudio5 qPCR-System (cat. # A28140, Applied Biosystems). Normalization was performed using two housekeeping genes, Hypoxanthine-Guanine Phosphoribosyltransferase (*Hprt*) and TATA Box Binding Protein (*Tbp*) combined. The following primer sequences were used: *Sp7*, forward GCCCCCTGGTGTTCTTCATT, reverse CCCATTGGACTTCCCCCTTC; *Bglap*, forward CCTGGCTGCGCTCTGTCT, reverse TGCTTGGACATGAAGGCTTTG; *Bmp2*, forward AGACCACCGGCTGGAGAG, reverse TTTCCCACTCATCTCTGGAAG; *Col1a1*, forward TGTTCAGCTTTGTGGACCTC, reverse TCAAGCATACCTCGGGTTTC; *Hprt*, forward GGTTAAGCAGTACAGCCCCA, reverse GGCCTGTATCCAACACTTCG; *Tbp*, forward CCAGAACAACAGCCTTCCAC, reverse GTGGAGTAAGTCCTGTGCCG.

### RNA sequencing

Hydrogels (viscous) with encapsulated cells were collected at directly before the application of induction medium and at different time points of the differentiation process (2h, 12h, 144h). For each time point, two samples from two independent experiments were used for each condition (ISO, HYPO, ISO_mech+_). Gels were chopped using a scalpel and subsequently transferred into DNAse free tubes, which were immediately frozen in liquid nitrogen. RNA isolation from samples was performed using the RNeasy Micro Kit (cat. # 74004, QIAGEN) according to the manufacturer’s protocol. mRNA-Seq libraries were prepared using KAPA mRNA HyperPrep Kit (cat. # 08098123702, Roche) according to the manufacturer’s protocol. RNA quality was checked using Bioanalyzer RNA kit (Agilent). Samples with RIN score >7 were used in the downstream library preparation process. In short, libraries were constructed from 300 ng of intact RNA. Size distribution and concentration of libraries were confirmed with Bioanalyzer and qPCR.

RNA-seq reads were quantified using Salmon v1.5.2^74^ using the GENCODE Release M28 (GRCm39) as the reference transcriptome^75^. Transcript quantification was performed in mapping-based mode with GC bias correction and additional inferential replicates with the parameters ‘--gcBias --numGibbsSamples 20 --thinningFactor 100’. Quantifications and corresponding sample metadata were imported with tximeta v1.10.0^76^ into R v4.1.0 for downstream analysis. Exploratory analysis and consistency between replicates were assessed by principal component (PCA) analysis on variance stabilizing transformed (VST) counts. PCA visualization was done on the top 250 genes with highest variance. For differential analysis with DESeq2 v1.32.0^77^, we used a design formula which models the stimulus-specific differences over time, accounting for baseline at time 0h (directly before induction) and batch variations. A gene was considered differentially expressed, (i.e., DEG), when its absolute L2FC value was ≥ 0.5 and its FDR-adjusted p-value was ≤ 0.1. To define specific modules of gene expressions during differentiation, we clustered DEGs based on their change over time by k-clustering (PAM k-medoids = 4) as implemented by R package cluster v2.1.2^78^. Gene Ontology (GO) enrichment analysis on DEG clusters was done using package clusterProfiler v4.0.0^79^. For enrichment analysis, the set of background genes was defined as all genes which were detectably expressed (i.e., count ≥ 10) at baseline 0h.

### Statistical analysis and illustration

The employed statistical tests are noted with the figures. Generally, to compare two groups, two-tailed unpaired t test was performed, and to compare multiple groups, one-way ANOVA with Tukey correction was performed. If standard deviations were significantly different between compared groups, the Brown-Forsythe ANOVA was used. Correlation analysis by Spearman’s rank correlation was used for mechanics (*E*, *τ*_1⁄2_) of viscous hydrogels, Brillouin shift, and nucleus staining heterogeneity with incubation osmolality. Statistical analysis was performed with GraphPad Prism (version 10.6.1). Illustrations in Fig. 1b and Fig. 5k were created with Biorender.com.

## Acknowledgements

We thank Dag Wulsten, Gao Xiang (Charité Berlin) for technical assistance with biomechanical characterization; Simon Reinke, Antje Blankenstein, Johanna Penzlin (Charité Berlin) for the provision of human bone fracture haematoma samples; Janosch Schoon and Sven Geissler for the provision of samples of bone marrow plasma and cerebrospinal fluid (Charité Berlin); Mario Thiele for technical assistance with microscopy image analysis (Charité Berlin); Norma Schulz, Aysha Bonell for technical assistance with RNA isolation and qPCR measurements (Charité Berlin); Maximilian Ebisch for performing and analyzing DSC measurements (BAM); Fabian Kriegel for performing and analyzing ICP-MS characterization (German Federal Institute for Risk Assessment); Aline Lückgen, Julia Löffler, Christian Bucher, Enrico Klotzsch, Clemens Franke (Charité Berlin), Georgios Kotsaris (Freie Universität Berlin), Dilara Perver, Fabrizio Pennacchio (ETH Zurich) for sharing their insights and expertise within the context of this work. This work was supported by the European Research Council (ERC-2021-ADG, 101054501), by the Einstein Foundation Berlin, and by the German Research Foundation (CRC 1444).

## Data availability

The main data supporting the results in this study are available within the paper and its Supplementary Information. The raw and analysed datasets generated during the study are available for research purposes from the corresponding author on reasonable request.

## Supplementary Information

**Supplementary Fig. 1.**
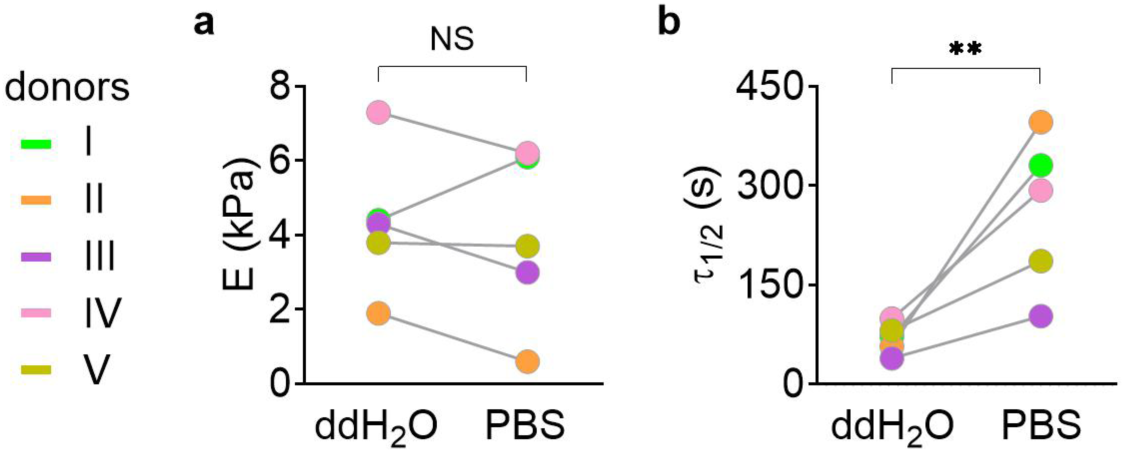
Mechanical characterization results of ex vivo human fracture haematoma for individual donors. Tissue samples from five donors (I–V) were characterized using unconfined uniaxial compression testing. A fresh, untreated tissue sample was dissected into similar sized samples of adequate size and form for mechanical characterization by uniaxial compression. Subsequently, dissected tissue samples of a donor were incubated in solutions for an extreme comparison of osmotic concentrations using ddH_2_O (hypo-osmolar) or PBS (iso-osmolar) to analyze effects of an osmolality decrease on viscoelastic properties (*E*, *τ*_1⁄2_). (**a**) *E* of fracture haematoma did not differ significantly between the two incubation conditions (*n* = 5 donors, NS *P* > 0.05 by Student’s t test). Comparing *E* of samples from individual donors after incubation showed that in four out of five donors patients increased values in hypo-osmolar compared to iso-osmolar condition; *E* increased at different extents (3%, 18%, 43%, 216%). For one donor, *E* decreased (28%) in hypo-osmolar compared to iso-osmolar condition instead. A larger cohort study would be required to clarify this aspect. Increased *E* in hypo-osmolar (i.e., low ion concentration) solutions was previously reported for cartilage^3,9^. (**b**) *τ*_1⁄2_ of fracture haematoma was decreased significantly in hypo-osmolar compared to iso-osmolar condition (*n* = 5 donors, \*\**P* < 0.01 by Student’s t test). Comparing *τ*_1⁄2_ of tissue samples of individual donors after incubation showed for all five donors lower values in hypo-osmolar compared to iso-osmolar. This showed that decreased environmental osmolality had a robust and stronger effect on *τ*_1⁄2_, enhancing viscous (i.e., stress relaxing) properties of fracture haematoma samples.

**Supplementary Fig. 2.**
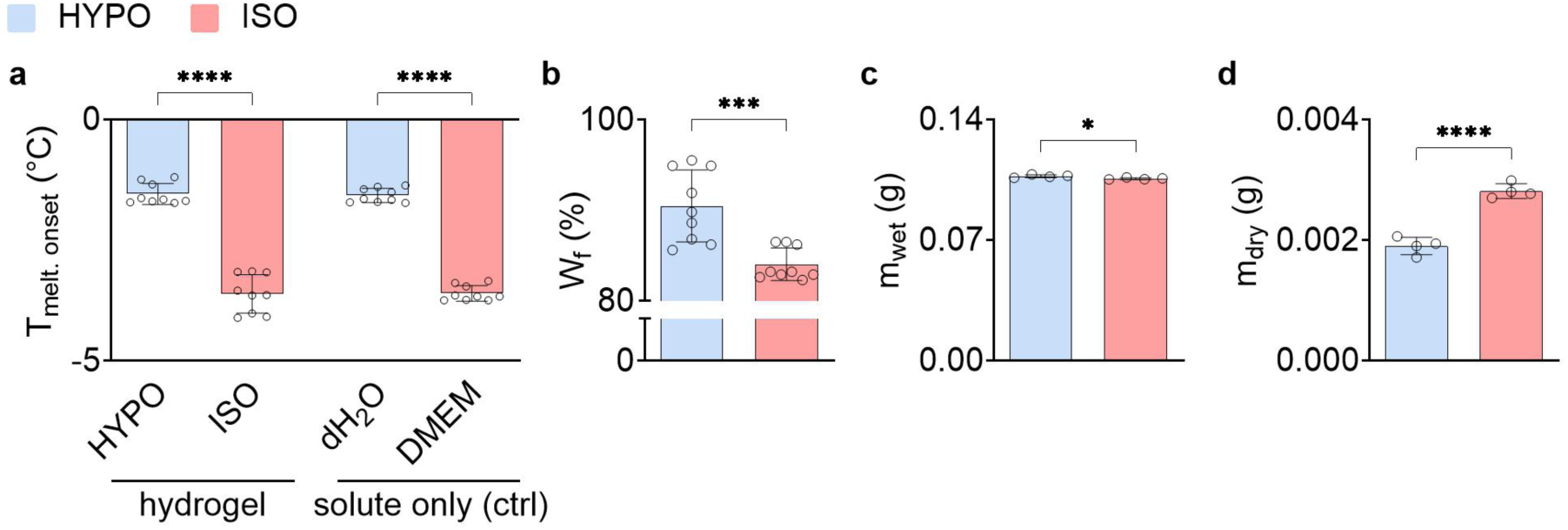
HYPO compared to ISO environment increases freezable water content in hydrogels, likely due to ion diffusion. (**a**) Melting onset temperature (T_melt_), quantified by DSC, of hydrogels after incubation in HYPO or ISO environment compared to the respective solute only as controls (i.e., ddH_2_O and DMEM for HYPO and ISO, respectively). The similar differences HYPO and ISO for both incubated hydrogels and the incubation solutions underline the effect of the osmotic environment modulation on the water binding properties by the altered ion concentration. This is in line with the high water content in the hydrogels produced with just 2wt% alginate (*n* = 9 samples, \*\*\*\**P* < 0.0001 by Student’s t test). (**b**) Analysis of the freezable water content (W_f_) using DSC showed increased W_f_ of hydrogels in HYPO compared to ISO, suggesting altered water binding (*n* = 9 samples, \*\*\**P* < 0.001 by Student’s t test). These results for alginate hydrogels were in line with previous observations for PEG hydrogels^80^. (**c**) Weight measurements showed only a very small increase of ∼1% in wet weight in HYPO (ca. 0.107 g) compared to ISO (0.106 g) (*n* = 4 samples, \**P* < 0.05 by Student’s t test). The measured weights were in line with the produced cylindrical hydrogels of 8 mm diameter and 2 mm height (volume 0.1005 cm^3^) considering the density of water (1.0 g/cm^3^) and the high water content in the hydrogel mix with just 2wt% alginate. The very small swelling difference in the extreme comparison (ddH_2_O versus DMEM) analyzed here suggested that likely there is no relevant change in cell swelling between the osmotic conditions (focus range) used for cell culture. There, the difference in osmolality between HYPO (250 mOsm/kg) and ISO (320 mOsm/kg) was ∼78% smaller than in the extreme comparison (ddH_2_O versus DMEM) analyzed here. (**d**) The quantification of dry weight showed a decrease in in HYPO (ddH_2_O) compared to ISO (DMEM) (*n* = 4 samples, \*\*\*\**P* < 0.0001 by Student’s t test). The hydrogels had first been equilibrated in ISO buffer after production and then been subjected to the different osmotic conditions. This suggested ion efflux from the hydrogels and would be expected due to the decrease in environmental osmolality for gels exposed to ddH_2_O. Further, this is in line with the measured T_melt_ values for incubated hydrogels and incubation solutes only, as a higher ion concentration would be expected to lower T_melt_. (*n* = 4 samples, \*\*\*\**P* < 0.0001 by Student’s t test). The data suggests that in HYPO compared to ISO condition, the negative charges of alginate chains retain more water molecules instead of ions that diffused out of the hydrogels. Bar plots: mean ± SD.

**Supplementary Fig. 3.**
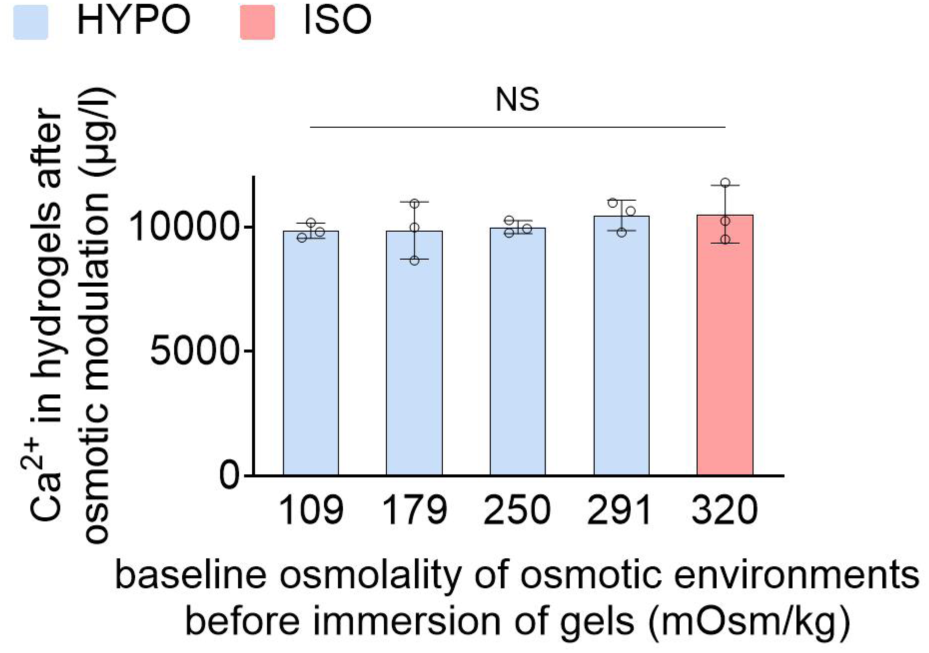
Calcium concentration inside hydrogels is not altered after one day of exposure to different hypo-osmolar environments, following one day of equilibration of ISO medium. Inductively coupled plasma mass spectrometry (ICP-MS) was used to quantify calcium concentration inside hydrogels. Ionically (calcium) crosslinked hydrogels were disintegrated using EDTA in PBS (without calcium and magnesium). Potential influence of the solution used for disintegrating the gels was considered in the calculations of calcium concentrations inside gels. Gels were produced in one batch (*n* = 3 samples, NS *P* > 0.05 by one-way ANOVA). Bar plots: mean ± SD.

**Supplementary Fig. 4.**
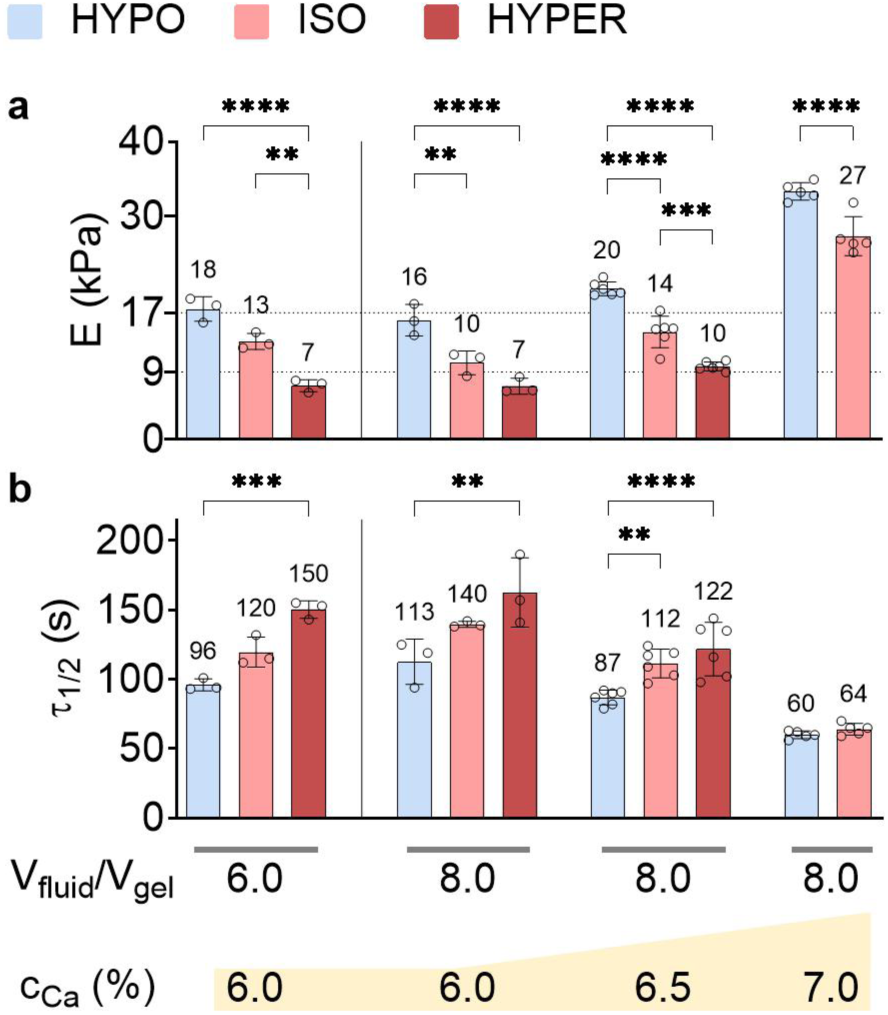
Environment osmolality and volume ratio of environment fluid and gel tune viscoelastic properties of ionically crosslinked alginate hydrogels. HYPO, ISO, and HYPER conditions refer to the osmotic concentrations 250, 320, and 390 mOsm/kg, respectively. The volume ratio of environment fluid and gel (V_fluid_/V_gel_) was 6.0 or 8.0. Ionic (calcium) crosslinking concentration (c_Ca_) were 6.0, 6.5, or 7.0%. Mechanical properties were quantified using uniaxial compression testing after one day of equilibration in ISO medium after hydrogel casting and subsequently, one day of osmotic modulation with HYPO, ISO, or HYPER environment. (**a**) Elastic modulus, E, and (**b**) stress relaxation half time, *τ*_1⁄2_ of hydrogels (*n* = 3–6 samples, \*\*\*\**P* < 0.0001, \*\*\**P* < 0.001, and \*\**P* < 0.01 by one-way ANOVA). The gels were produced in one batch. The gels used for investigations on cell-matrix interactions in the focus range of osmotic environments (250, 320, 390 mOsm/kg) were produced at c_Ca_ =6.5% and incubated at V_fluid_/V_gel_ =8.0. The gels used to investigate the influence of enhanced viscoelasticity at the iso-osmolar (ISO, 320 mOsm/kg), referred to as ISO_mech+_ condition, were produced at c_Ca_ =7.0% and incubated at V_fluid_/V_gel_ =8.0. Bar plots: mean ± SD.

**Supplementary Fig. 5.**
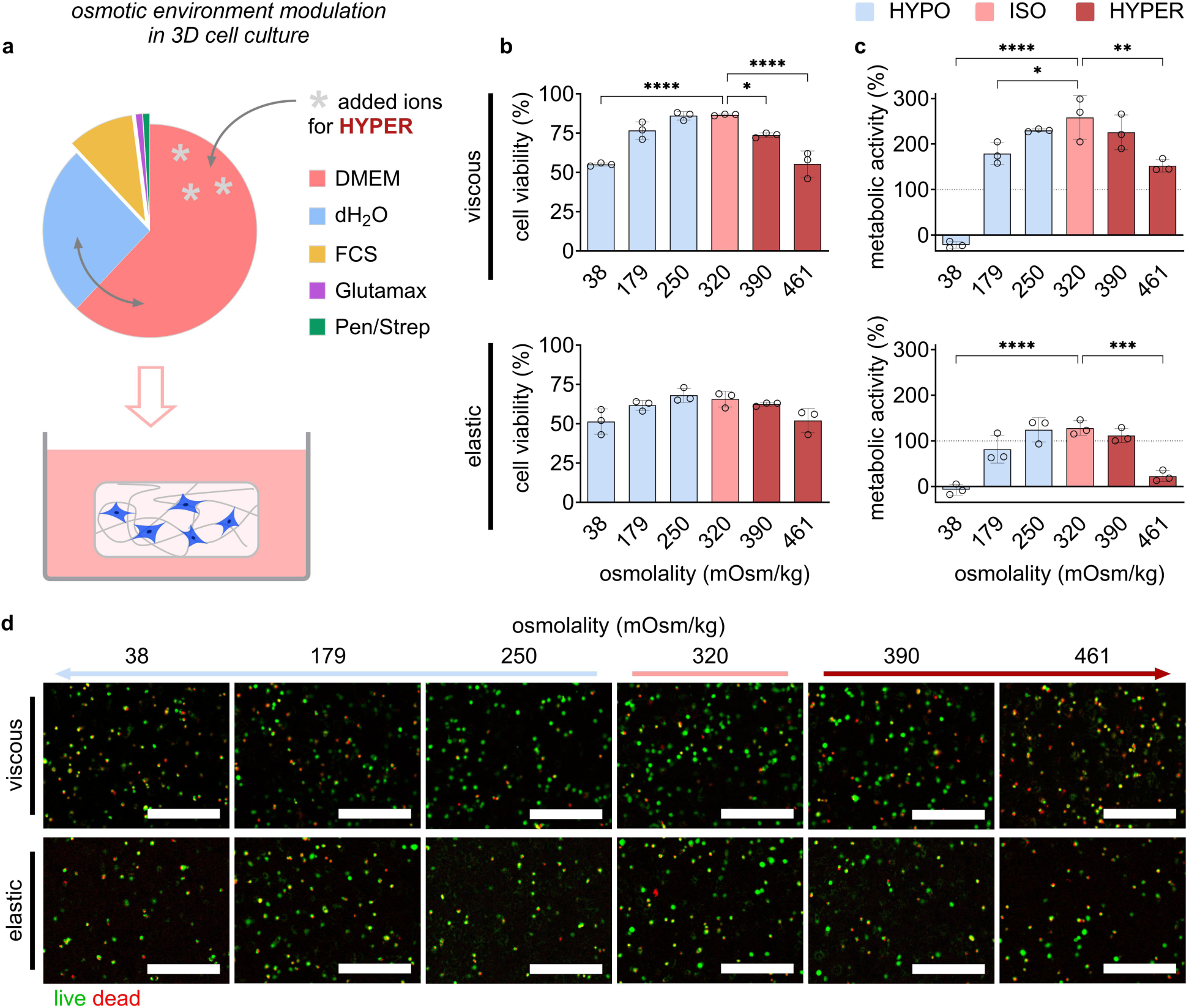
Modulating environmental osmolality while maintaining cell viability to mimic the osmolality drop in 3D culture. A focus range of osmotic concentrations for maintained MSC viability and metabolic activity in hydrogels was established based on quantifications of cell viability from live/dead staining and metabolic activity by alamarBlue assay. (**a**) Illustration of the experimental setup for modulating the environmental osmolality for 3D cell culture in hydrogels using the defined cell culture medium formulation. The medium formulation consisted of Dulbecco’s Modified Eagle Medium (DMEM), deionized water (dH_2_O), fetal calf serum (FCS), GlutaMAX supplement (Glutamax), and Penicillin-Streptomycin (Pen/Strep). The osmotic concentrations of tested incubation conditions were for HYPO, 38, 179, and 250 mOsm/kg, for ISO, 320 mOsm/kg, and for HYPER, 390 and 461 mOsm/kg. Quantification of (top: viscous; bottom: elastic) (**b**) cell viability (live/dead staining) and (**c**) cell metabolic activity (alamarBlue assay) after two days of culture in viscous or elastic gels in a wide range of osmotic concentrations (*n* = 3 samples per condition, \*\*\*\**P* < 0.0001, \*\*\**P* < 0.001, \*\**P* < 0.01, and \**P* < 0.05 by one-way ANOVA, comparisons with ISO are shown). (**d**) Representative microscopy images of live (green) / dead (red) staining from which cell viability was quantified in viscous (top) or elastic (bottom) gels in different osmotic concentrations. Scale bar = 100 µm; bar plots: mean ± SD.

**Supplementary Fig. 6.**
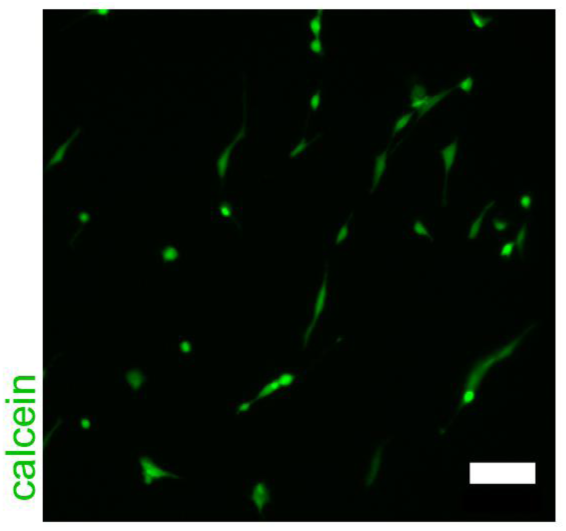
Calcein staining of MSCs seeded on top of covalently UV crosslinked hydrogels (elastic) show cell adhesion and confirm successful coupling of RGD ligands. Covalently crosslinked hydrogels were produced as described but without encapsulation of cells. MSCs were seeded on top of the hydrogels and allowed to attach during incubation in iso-osmolar expansion medium overnight. Calcein staining was used to stain the cell body of live cells, similar to the method described for live dead staining. Live cell microscopy was used to image cells. The provided representative image shows cells spreading to elongated shape, which confirms adhesion of the cells to the hydrogel. Scale bar: 100 µm.

### Supplementary Note 1 Mesh size analysis of hydrogels using SAXS

SAXS measurements were performed from three ionically crosslinked hydrogels (viscous) after incubation in ddH_2_O (0 mOsm/kg), HYPO (250 mOsm/kg), or ISO (320 mOsm/kg) solution and without further samples treatment. The resultant scattering curves are provided in Supplementary Fig. 7 (black solid lines). The scattering intensity of the samples characteristically decays by about five orders of magnitude in the available *q*-range of 0.08 nm^-1^ to 7.0 nm^-1^. This *q*-range corresponds to a length scale of about 40 nm to 0.5 nm. Therefore, the data provide information on the structure of the gel in this size range. Schematically, the structure of alginate is typically interpreted as a mesh of rods of primary and secondary rods (see for example Posbeyikian et al.^81^). The primary rods are formed by dimers of alginate chains and the secondary rods are formed by aggregated primary rods. These rod-like structures are considered relatively stiff in the junction zones and are known as the “broken rod” model^82–84^.

The structural features of such a gel network build-up from thinner and thicker rods contribute differently to the scattering intensity at different *q*-values of the SAXS patterns. We found that the overall scattering intensity *I(q)* can suitably be modeled as

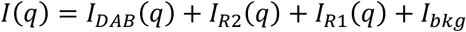

The first summand represents the mesh of the gel in terms of the Debye-Anderson-Brumberger model^85^

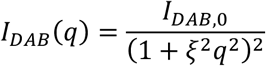

where *I_DAB_*_,0_is the scattering of *I_DAB_*(*q*) at *q*=0 and *ξ* is the correlation length of the mesh. Descriptively, *ξ* can be interpreted as mesh size.

The scattering of a rodlike segment with a radius *R_i_* and a length *L_i_* is given as^86,87^

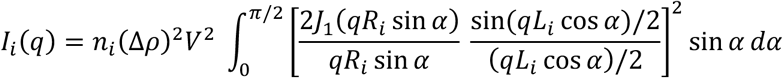

where *n_i_* is the number concentration of the rods, Δ*ρ* is the difference in scattering length density between the rods and the buffer, *V* is the volume of a rod, and *J*_1_ is a first order Bessel function. Finally, a remaining scattering background of low intensity *I_bk_*_g_ - originating from density fluctuations - is taken as constant.

Best fits of the data employing this model are provided in Supplementary Fig. 7. It can be seen there that the general shape of the data is well represented by this model (red solid curves). The respective individual contribution of *I_DAB_*(*q*), *I_R_*_2_(*q*), *I_R_*_1_(*q*) and *I_bk_*_g_ are also displayed for comparison. The lengths of the primary and secondary filaments were both held constant at a value of 200 nm to avoid ambiguous fit results.

The primary filaments radii were determined as *R*_1_ = 0.6 nm for the three samples. It seems reasonable to obtain the same *R*_1_-value for the three samples, if we assume that this represents the cross-sectional radius of an alginate dimer. Depending on the alginate sample type and preparation conditions, cross-sectional radii in the range of 0.54 nm and 2.22 nm have been reported^84^. But most of their values are around 0.6 nm. Similarly, values for the cross-section radius were reported also around 0.6 nm^81^. Therefore, we can conclude that the value that we obtained for cross-section radius for the primary filament agrees with the literature.

The cross-section radii for the secondary filaments were determined as 6.0 nm (ddH_2_O), 3.2 nm (HYPO), and 1.8 nm (ISO). This indicates a trend that *R*_2_ decreases with increasing osmolality at sample preparation. Previously, values for *R*_2_ were reported in the range of about 1.5 nm to 3.8 nm^84^ and around 2.5 nm^81^. These literature values are in agreement with our data. Our data suggest that cross-section radii for the secondary filaments were comparable in HYPO and ISO conditions (3.2 nm and 0.8 nm), but clearly larger in ddH_2_O condition (6.0 nm).

The mesh-size parameter *ξ* of was determined as 19.0 nm (ddH_2_O), 8.5 nm (HYPO), and 9.7 nm (ISO), showing *ξ* in ddH_2_O was much larger than in HYPO and ISO conditions. This suggests that the mesh of a hydrogel in ddH_2_O condition was much more open than that of a hydrogel in HYPO or ISO condition, while the mesh sizes of hydrogels in HYPO or ISO condition were very similar.

Tentatively, we estimated the ratio of primary to secondary filaments under the assumptions that the scattering contrast and the length are similar for both. The number ratios *N*_1_/*N*_2_ were determined as 232 (ddH_2_O), 59 (HYPO), and 7 (ISO). This suggests that the number of primary filaments outweighed the number of secondary filaments in all three cases. Furthermore, the ratio of primary to secondary filaments decreased with increasing osmolality.

In conclusion, we were able to characterize the alginate hydrogels with SAXS by utilizing a simplified structure model. SAXS data and model curves are summarized in Supplementary Table **1**. The mesh size of hydrogels in HYPO or ISO condition was similar (8.5 nm and 9.7 nm), while it was much larger for the extremely hypo-osmolar ddH_2_O condition (19.0 nm). This shows that the effect of the difference between HYPO and ISO conditions on the mesh size of the viscous hydrogels was negligible.

**Supplementary Table 1.** SAXS data and model curves (black and red solid lines, respectively) of hydrogels after incubation in ddH_2_O, HYPO, or ISO condition. Components of the model are the Debye-Anderson-Brumberger expression *I_DAB_*(*q*) for the mesh structure (dash-dotted blue curves), rods for the secondary *I_R_*_2_(*q*) and primary filaments *I_R_*_1_(*q*) (dashed magenta and dotted green curves) and constant background intensity *I_bk_*_g_ (dashed black lines).

| **condition** | ***ξ*** | ***H*** | ***N***<sub>**1**</sub> | ***R***<sub>**1**</sub> | ***N***<sub>**2**</sub> | ***R***<sub>**2**</sub> | ***N***<sub>**1**</sub> **/*N***<sub>**2**</sub> |
|---|---|---|---|---|---|---|---|
| ddH<sub>2</sub>O | 19.0 | 0.5 | 2E-07 | 0.6 | 8.61E-10 | 6.0 | 232 |
| HYPO | 8.5 | 0.5 | 6E-08 | 0.6 | 1.01E-09 | 3.2 | 59 |
| ISO | 9.7 | 0.5 | 3E-08 | 0.6 | 4.1E-09 | 1.8 | 7 |

**Supplementary Fig. 7.**
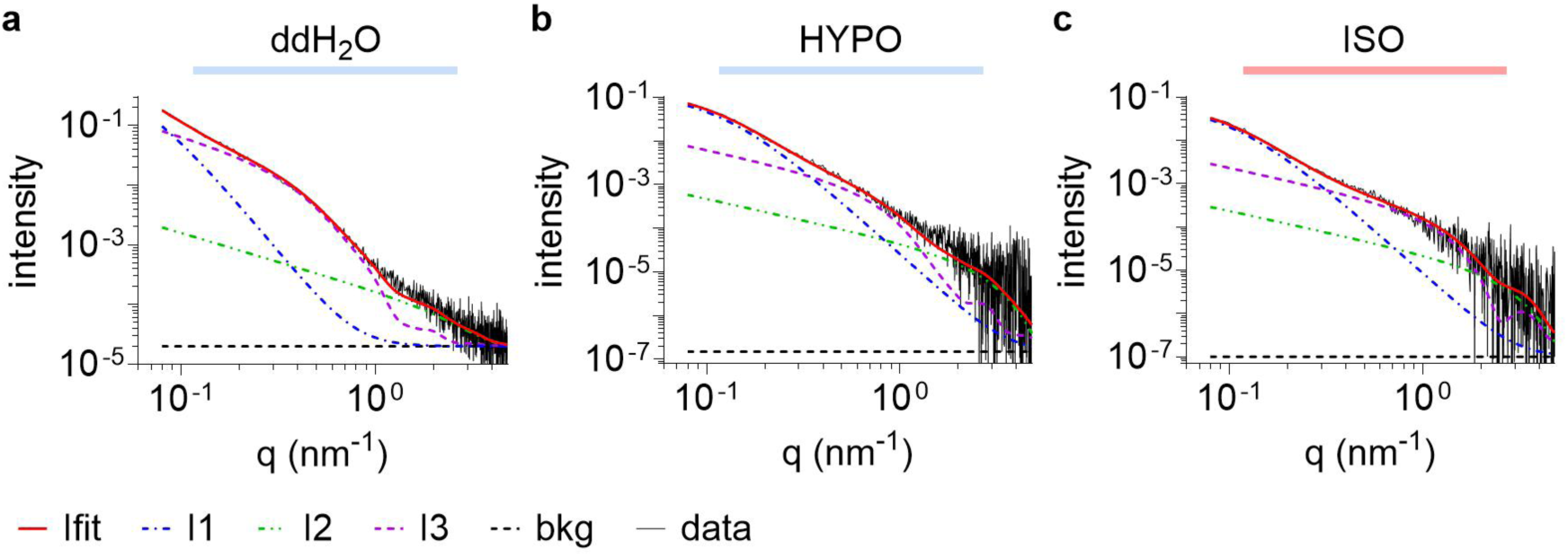
SAXS analysis shows no change in hydrogel mesh size between HYPO and ISO condition, but in ddH_2_O compared to HYPO and ISO. SAXS measurements were performed from samples of ionically crosslinked hydrogels (viscous) after incubation in (**a**) ddH_2_O (0 mOsm/kg), (**b**) HYPO (250 mOsm/kg), or (**c**) ISO (320 mOsm/kg) solution. Scattering curves (black solid lines) and model curves (red solid lines) are displayed. Contributions of the components of the model are the Debye-Anderson-Brumberger mesh model (blue dashed-dotted lines), the secondary filaments (magenta dashed lines), the primary filaments (green dashed and double dotted lines) and a low constant background (black dashed lines). Hydrogel mesh size was compared between HYPO (250 mOsm/kg) and ISO (320 mOsm/kg), and with respect to an extremely hypo-osmolar condition (ddH_2_O, 0 mOsm/kg). While a decrease in mesh size was observed for hydrogels incubated in ddH_2_O compared to HYPO and ISO conditions, no difference was found between HYPO and ISO conditions. This shows that the change in osmolality between HYPO and ISO conditions did not affect the mesh size of the incubated hydrogels.

**Supplementary Fig. 8.**
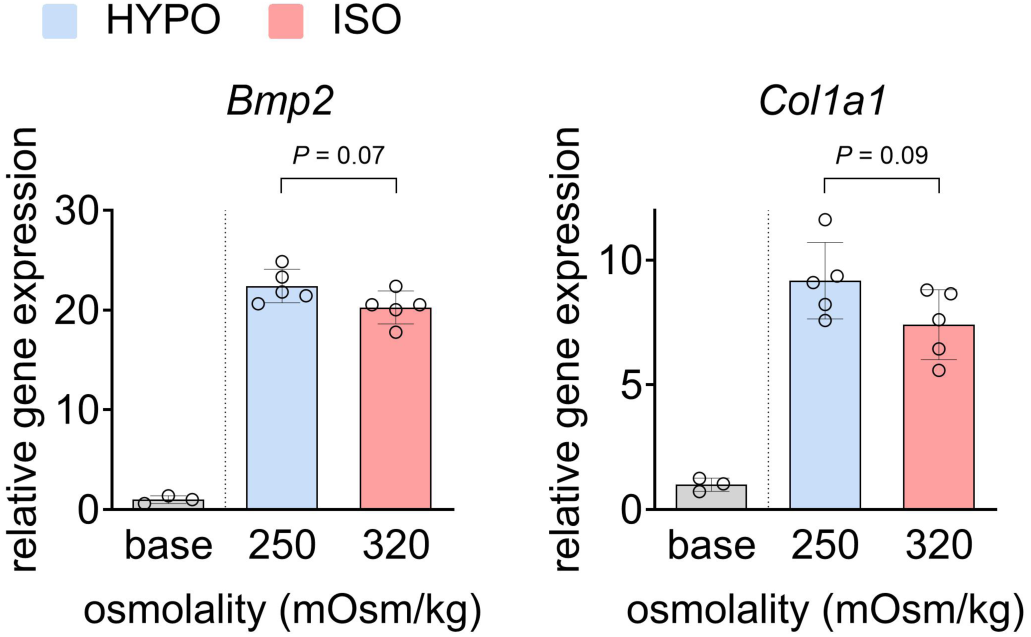
Relative gene expression of osteogenic genes analyzed by qPCR. Analysis of gene expression of *Bmp2* and *Col1a1*, after four days of osteogenic induction in viscous ECM in HYPO compared to ISO conditions. Base conditions represent cells harvested after equilibration in hydrogels, right before the start of incubation with osteogenic induction medium. Bar plots: mean ± SD.

**Supplementary Fig. 9.**
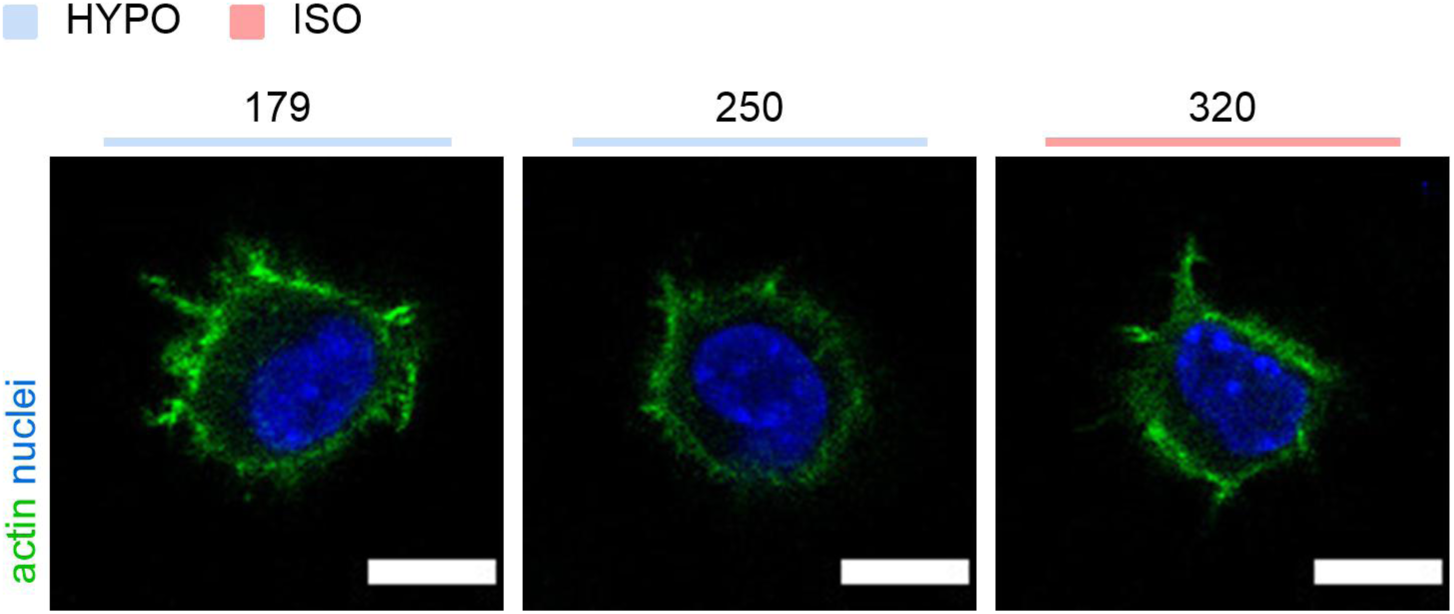
Osmolality drop causes immediate change in nucleus homogeneity suggesting chromatin decondensation. Nucleus (DAPI) and actin (Phalloidin) staining were used to analyze for changes in chromatin structure caused by modulation of environmental osmolality. Representative images of staining in 179, 250, and 320 mOsm/kg conditions are shown. Scale bars: 10 µm.

**Supplementary Fig. 10.**
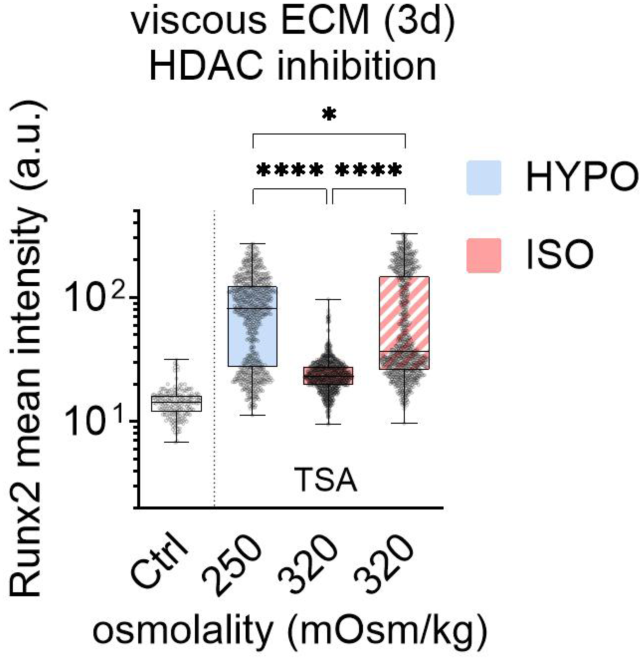
Quantification of Runx2 TF expression. Nuclear mean staining of Runx2 after three days of osteogenic induction in viscous ECM hydrogels to assess the influence of chromatin decompaction on MSC osteogenic differentiation using HDAC inhibition by TSA. Comparison of the three conditions ISO supplemented with 75 nM TSA, HYPO, and standard ISO (*n* = 750–839 single cells per condition, \*\*\*\**P* < 0.0001, and \**P* < 0.05 by Kruskal-Wallis test).

### Supplementary Note 2 Effect of osmotic pressure on MSC osteogenic differentiation

Previous work investigated cell volume as a regulator of MSC osteogenic differentiation in 3D viscoelastic hydrogels by supplementing culture medium with PEG to constrict cell volume, which decreased osteogenic differentiation^35^. In the same study, dilution of the culture medium from day two of osteogenic induction was used to increase cell volume and enhanced osteogenic differentiation, generally consistent with our observation that hypo-osmolality promotes osteogenic differentiation.

We asked whether the pro-regenerative effect of the osmolality drop observed in our study could be explained primarily by persistent cell volume expansion. To test this, we added PEG to HYPO medium to compensate the reduced osmolality by macromolecular osmotic pressure while maintaining the lowered ionic concentration. If persistent volume expansion were the dominant mechanism in our experiments, this compensation condition would be expected to restore differentiation toward ISO levels. In contrast, applying 70 mOsm/kg PEG to HYPO medium (250+70PEG) strongly reduced ALP and actin staining compared with both HYPO (250 mOsm/kg) and ISO (320 mOsm/kg) conditions (Supplementary Fig. 11a– b).

We further assessed whether PEG altered the hydrogel mechanics or impaired cell viability. Adding PEG to hypo-osmolar buffer caused only a minor change in elastic modulus and did not alter stress relaxation half-time (Supplementary Fig. 11c–d). Addition of 70 mOsm/kg PEG did not reduce cell viability or metabolic activity, whereas a higher PEG concentration reduced viability (Supplementary Fig. 11e–f). Thus, the reduced osteogenic differentiation in the PEG-compensated condition was unlikely to result from altered hydrogel stress relaxation, impaired viability, or reduced metabolic activity.

Together, these data indicate that the osmolality-drop stimulus identified here is mechanistically distinct from persistent cell volume modulation by macromolecular osmotic pressure. Our injury-inspired protocol applies a hypo-osmolar shift at the onset of osteogenic induction, maintains constant supplement concentrations, and accounts for ECM mechanics. This design revealed a regenerative stimulus driven by coupled osmotic and mechanical changes in the extracellular microenvironment.

**Supplementary Fig. 11.**
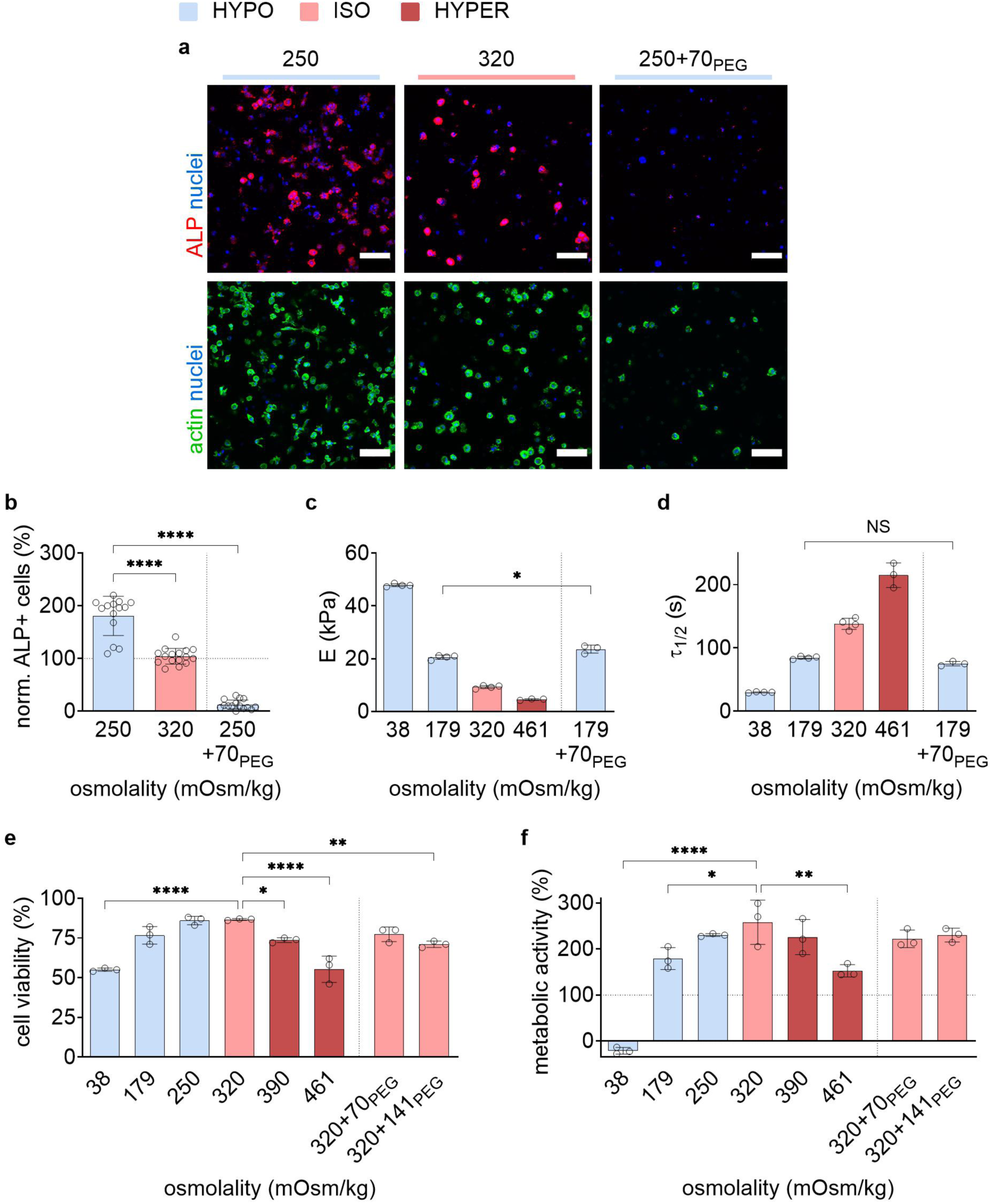
Compensation of change in environmental osmolality due to osmolality drop by addition of PEG to culture medium does not restore osteogenic differentiation performance of the reference condition ISO but leads to strong decrease. Osmotic pressure was applied in media of different ionic concentrations (i.e., 179, 250, 320) by addition of polyethylene glycol (PEG). The osmotic concentration (in mOsm/kg) of PEG added to a condition was depicted by +70PEG or +141PEG. PEG of low molecular weight (PEG-400, Mw=400 Da) was used to diffuse into hydrogels and apply osmotic pressure to encapsulated cells (as PEG would not diffuse into cells as ions could), similar to the method used by Lee et al. (2019) ^35^. The conditions 250 and 250+70PEG had the same ionic concentration (i.e., HYPO medium), while 250+70PEG had 70 mOsm/kg of PEG added to equal 320 mOsm/kg overall osmotic concentration. The reference iso-osmolar condition (320) consisted of ISO medium with higher ionic concentration than 250 and 250+70PEG, but the same overall osmotic concentration as 250+70PEG. Osteogenic differentiation was evaluated by alkaline phosphatase (ALP) staining after six days of induction at different osmotic conditions in the viscous hydrogels. (**a**) Representative images of nuclei and ALP staining (top) and nuclei and actin staining (bottom). (**b**) Quantification of ALP staining by ALP+ cells normalized by the ISO condition (*n* = 14–16 images from three samples per condition, \*\*\*\**P* < 0.0001 by one-way ANOVA, only comparisons to 250 mOsm/kg condition are shown). To assess to what extent hydrogel mechanics could be affected by addition of 70 mOsm/kg of PEG-400 to hypo-osmolar culture media, mechanical characterization was performed for one hypo-osmolar condition as an example (179 mOsm/kg). (**c**) Elastic modulus (*E*) and (**d**) stress relaxation half time (*τ*_1⁄2_) were quantified (*n* = 3–4 samples per condition, \**P* < 0.05, and NS *P* > 0.05 by one-way ANOVA, only the comparison between 179 and 179+70PEG conditions is shown). No differences in stress relaxation behavior, quantified was observed. The slight increase in *E* was considered negligible for the outcome of the performed experiment as the investigated range of stiffness (e.g., HYPO: 20 kPa, ISO: 14 kPa) was previously shown to favor osteogenic differentiation in similar viscous gels^14^. In this range of viscoelastic properties, enhanced ECM stiffness or stress relaxation would be expected to increase osteogenic differentiation, rendering them unlikely to be the cause for the strong decrease in osteogenic differentiation in the 250+70PEG condition. Quantification of (**e**) cell viability and (**f**) metabolic activity of MSCs showed neither was not affected by the addition of 70 mOsm/kg. Thus, also impaired viability or metabolic activity were unlikely to cause the decreased osteogenic differentiation performance under applied osmotic pressure (*n* = 3 samples per condition, \*\*\*\**P* < 0.0001, \*\**P* < 0.01, and \**P* < 0.05 by one-way ANOVA, only comparisons to 320 mOsm/kg (ISO) condition are shown). Bar plots: mean ± SD.

## References

1. Knepper, M. A., Saidel, G. M., Hascall, V. C. & Dwyer, T. Concentration of solutes in the renal inner medulla: interstitial hyaluronan as a mechano-osmotic transducer. Am. J. Physiol. Physiol. 284, F433–F446 (2003).

2. Buschmann, M. D. & Grodzinsky, A. J. A molecular model of proteoglycan-associated electrostatic forces in cartilage mechanics. J. Biomech. Eng. 117, 179–192 (1995).

3. Mow, V. C., Ratcliffe, A. & Robin Poole, A. Cartilage and diarthrodial joints as paradigms for hierarchical materials and structures. Biomaterials 13, 67–97 (1992).

4. Wiig, H. Pathophysiology of tissue fluid accumulation in inflammation. J. Physiol. 589, 2945–2953 (2011).

5. Soll, D., Chu, C., Sun, S., Lutz, V., Arunkumar, M., Gachechiladze, M., Schäuble, S., Alissa-alkhalaf, M., Nguyen, T., Khalil, M., Garcia-ribelles, I., Mueller, M., Buder, K., Michalke, B., Moosmann, C., Schamel, W. W., Huber, M. & Zielinski, C. E. Sodium chloride in the tumor microenvironment enhances T cell metabolic fitness and cytotoxicity. Nat. Immunol. 25, 1830–1844 (2024).

6. Nia, H. T., Munn, L. L. & Jain, R. K. Physical traits of cancer. Science 370, eaaz0868 (2020).

7. Mu, Y., Du, Z., Xiao, L., Gao, W., Crawford, R. & Xiao, Y. The Localized Ionic Microenvironment in Bone Modelling/Remodelling: A Potential Guide for the Design of Biomaterials for Bone Tissue Engineering. J. Funct. Biomater. 14, (2023).

8. Yeh, S. C. A., Hou, J., Wu, J. W., Yu, S., Zhang, Y., Belfield, K. D., Camargo, F. D. & Lin, C. P. Quantification of bone marrow interstitial pH and calcium concentration by intravital ratiometric imaging. Nat. Commun. 13, 1–13 (2022).

9. Eisenberg, S. R. & Grodzinsky, A. J. Swelling of articular cartilage and other connective tissues: Electromechanochemical forces. J. Orthop. Res. 3, 148–159 (1985).

10. Kollert, M. R., Krämer, M., Brisson, N. M., Schemenz, V., Tsitsilonis, S., Qazi, T. H., Fratzl, P., Vogel, V., Reichenbach, J. R. & Duda, G. N. Water and ions binding to extracellular matrix drives stress relaxation, aiding MRI detection of swelling-associated pathology. *Nat*. Biomed. Eng. 9, 772– 786 (2025).

11. Roffay, C., Molinard, G., Kim, K., Urbanska, M., Andrade, V., Barbarasa, V., Nowak, P., Mercier, V., García-calvo, J., Matile, S., Loewith, R., Echard, A., Guck, J., Lenz, M. & Roux, A. Passive coupling of membrane tension and cell volume during active response of cells to osmosis. Proc. Natl. Acad. Sci. U. S. A. 118, e2103228118 (2021).

12. Jentsch, T. J. VRACs and other ion channels and transporters in the regulation of cell volume and beyond. Nat. Rev. Mol. Cell Biol. 17, 293–307 (2016).

13. Ehrlich, S., Wolff, N., Schneiderman, R., Maroudas, A., Parker, K. H. & Winlove, C. P. The osmotic pressure of chondroitin sulphate solutions: Experimental measurements and theoretical analysis. Biorheology 35, 383–397 (1998).

14. Chaudhuri, O., Gu, L., Klumpers, D., Darnell, M., Bencherif, S. A., Weaver, J. C., Huebsch, N., Lee, H. P., Lippens, E., Duda, G. N. & Mooney, D. J. Hydrogels with tunable stress relaxation regulate stem cell fate and activity. Nat. Mater. 15, 326–334 (2016).

15. Guo, M., Pegoraro, A. F., Mao, A., Zhou, E. H., Arany, P. R., Han, Y., Burnette, D. T., Jensen, M. H., Kasza, K. E., Moore, J. R., Mackintosh, F. C., Fredberg, J. J., Mooney, D. J., Lippincott-Schwartz, J. & Weitz, D. A. Cell volume change through water efflux impacts cell stiffness and stem cell fate. Proc. Natl. Acad. Sci. U. S. A. 114, E8618–E8627 (2017).

16. Masic, A., Bertinetti, L., Schuetz, R., Chang, S. W., Metzger, T. H., Buehler, M. J. & Fratzl, P. Osmotic pressure induced tensile forces in tendon collagen. Nat. Commun. 6, 5942 (2015).

17. Dolega, M. E., Monnier, S., Brunel, B., Joanny, J.-F. F., Recho, P. & Cappello, G. Extra-cellular matrix in multicellular aggregates acts as a pressure sensor controlling cell proliferation and motility. Elife 10, e63258 (2021).

18. Werbner, B., Zhou, M., McMindes, N., Lee, A., Lee, M. & O’Connell, G. D. Saline-polyethylene glycol blends preserve in vitro annulus fibrosus hydration and mechanics: An experimental and finite-element analysis. J. Mech. Behav. Biomed. Mater. 125, 104951 (2021).

19. Engler, A. J., Sen, S., Sweeney, H. L. & Discher, D. E. Matrix elasticity directs stem cell lineage specification. Cell 126, 677–689 (2006).

20. Vogel, V. & Sheetz, M. Local force and geometry sensing regulate cell functions. Nat. Rev. Mol. Cell Biol. 7, 265–275 (2006).

21. Discher, D. E., Janmey, P. & Wang, Y. L. Tissue cells feel and respond to the stiffness of their substrate. Science 310, 1139–1143 (2005).

22. Trappmann, B., Gautrot, J. E., Connelly, J. T., Strange, D. G. T., Li, Y., Oyen, M. L., Cohen Stuart, M. A., Boehm, H., Li, B., Vogel, V., Spatz, J. P., Watt, F. M. & Huck, W. T. S. Extracellular-matrix tethering regulates stem-cell fate. Nat. Mater. 11, 642–649 (2012).

23. Vogel, V. Unraveling the Mechanobiology of Extracellular Matrix. Annu. Rev. Physiol. 80, 353– 387 (2018).

24. Chaudhuri, O., Cooper-White, J., Janmey, P. A., Mooney, D. J. & Shenoy, V. B. Effects of extracellular matrix viscoelasticity on cellular behaviour. Nature 584, 535–546 (2020).

25. Duda, G. N., Geissler, S., Checa, S., Tsitsilonis, S., Petersen, A. & Schmidt-Bleek, K. The decisive early phase of bone regeneration. Nat. Rev. Rheumatol. 19, 78–95 (2023).

26. Schell, H., Duda, G. N., Peters, A., Tsitsilonis, S., Johnson, K. A. & Schmidt-Bleek, K. The haematoma and its role in bone healing. J. Exp. Orthop. 4, 5 (2017).

27. Brauer, E., Klein, O., Neblich, G., Duda, G. N., Petersen, A., Lippens, E., Klein, O., Nebrich, G., Schreivogel, S., Korus, G., Duda, G. N. & Petersen, A. Collagen fibrils mechanically contribute to tissue contraction in an in vitro wound healing scenario. Adv. Sci. 6, 1801780 (2019).

28. Vining, K. H., Marneth, A. E., Adu-Berchie, K., Grolman, J. M., Tringides, C. M., Liu, Y., Wong, W. J., Pozdnyakova, O., Severgnini, M., Stafford, A., Duda, G. N., Hodi, F. S., Mullally, A., Wucherpfennig, K. W. & Mooney, D. J. Mechanical checkpoint regulates monocyte differentiation in fibrotic niches. Nat. Mater. 21, 939–950 (2022).

29. Dargaville, B. L. & Hutmacher, D. W. Water as the often neglected medium at the interface between materials and biology. Nat. Commun. 13, 4222 (2022).

30. Gonella, G., Backus, E. H. G., Nagata, Y., Bonthuis, D. J., Loche, P., Schlaich, A., Netz, R. R., Kühnle, A., McCrum, I. T., Koper, M. T. M., Wolf, M., Winter, B., Meijer, G., Campen, R. K. & Bonn, M. Water at charged interfaces. Nat. Rev. Chem. 5, 466–485 (2021).

31. Mackie, W., Noy, R. & Sellen, D. B. Solution properties of sodium alginate. Biopolymers 19, 1839–1860 (1980).

32. Zhang, H., Wang, H., Wang, J., Guo, R. & Zhang, Q. The effect of ionic strength on the viscosity of sodium alginate solution. Polym. Adv. Technol. 12, 740–745 (2001).

33. Kruck, B., Zimmermann, E. A., Damerow, S., Figge, C., Julien, C., Wulsten, D., Thiele, T., Martin, M., Hamdy, R., Reumann, M. K., Duda, G. N., Checa, S. & Willie, B. M. Sclerostin Neutralizing Antibody Treatment Enhances Bone Formation but Does Not Rescue Mechanically Induced Delayed Healing. J. Bone Miner. Res. 33, 1686–1697 (2018).

34. Huebsch, N., Arany, P. R., Mao, A. S., Shvartsman, D., Ali, O. A., Bencherif, S. A., Rivera-Feliciano, J. & Mooney, D. J. Harnessing traction-mediated manipulation of the cell/matrix interface to control stem-cell fate. Nat. Mater. 9, 518–526 (2010).

35. Lee, H. P., Stowers, R. & Chaudhuri, O. Volume expansion and TRPV4 activation regulate stem cell fate in three-dimensional microenvironments. Nat. Commun. 10, 529 (2019).

36. Loebel, C., Mauck, R. L. & Burdick, J. A. Local nascent protein deposition and remodelling guide mesenchymal stromal cell mechanosensing and fate in three-dimensional hydrogels. Nat. Mater. 18, 883–891 (2019).

37. Loeffler, J., Duda, G. N., Sass, F. A. & Dienelt, A. The Metabolic Microenvironment Steers Bone Tissue Regeneration. Trends Endocrinol. Metab. 29, 99–110 (2018).

38. Bernardo, M. E. & Fibbe, W. E. Review Mesenchymal Stromal Cells : Sensors and Switchers of Inflammation. Stem Cell 13, 392–402 (2013).

39. Compan, V., Baroja-Mazo, A., López-Castejón, G., Gomez, A. I., Martínez, C. M., Angosto, D., Montero, M. T., Herranz, A. S., Bazán, E., Reimers, D., Mulero, V. & Pelegrín, P. Cell Volume Regulation Modulates NLRP3 Inflammasome Activation. Immunity 37, 487–500 (2012).

40. Chessa, F., Mathow, D., Wang, S., Hielscher, T. & Atzberger, A. The renal microenvironment modifies dendritic cell phenotype. Kidney Int. 89, 82–94 (2016).

41. Ferrari, K. J., Scelfo, A., Jammula, S. G., Cuomo, A., Barozzi, I., Stützer, A., Fischle, W., Bonaldi, T. & Pasini, D. Polycomb-Dependent H3K27me1 and H3K27me2 Regulate Active Transcription and Enhancer Fidelity. Mol. Cell 53, 49–62 (2014).

42. Mattei, A. L., Bailly, N. & Meissner, A. DNA methylation: a historical perspective. Trends Genet. 38, 676–707 (2022).

43. Perino, M. & Veenstra, G. J. C. Chromatin Control of Developmental Dynamics and Plasticity. Dev. Cell 38, 610–620 (2016).

44. Irianto, J., Swift, J., Martins, R. P., McPhail, G. D., Knight, M. M., Discher, D. E. & Lee, D. A. Osmotic challenge drives rapid and reversible chromatin condensation in chondrocytes. Biophys. J. 104, 759–769 (2013).

45. Byfield, F. J., Eftekhari, B., Kaymak-loveless, K., Mandal, K., Li, D., Wells, R. G., Chen, W., Brujic, J., Bergamaschi, G., Wuite, G. J. L., Patteson, A. E. & Janmey, P. A. Metabolically intact nuclei are fluidized by the activity of the chromatin remodeling motor BRG1. Biophysj 124, 494– 507 (2025).

46. Stephens, A. D., Banigan, E. J., Adam, S. A., Goldman, R. D. & Marko, J. F. Chromatin and lamin A determine two different mechanical response regimes of the cell nucleus. Mol. Biol. Cell 28, 1984–1996 (2017).

47. Corato, M. De & Gomez-benito, M. J. Interplay of chromatin organization and mechanics of the cell nucleus. Biophysj 123, 3386–3396 (2024).

48. Carrivain, P., Cournac, A., Lavelle, C., Lesne, A., Mozziconacci, J., Paillusson, F., Signon, L., Victor, J. M. & Barbi, M. Electrostatics of DNA compaction in viruses, bacteria and eukaryotes: Functional insights and evolutionary perspective. Soft Matter 8, 9285–9301 (2012).

49. Bloomfield, V. A. DNA condensation by multivalent cations. Biopolymers 44, 269–282 (1997).

50. Paine, P. L., Moore, L. C. & Horowitz, S. B. Nuclear envelope permeability. Nature 254, 109–114 (1975).

51. Lima, A. F., May, G., Colunga, J., Pedreiro, S., Paiva, A., Ferreira, L., Enver, T., Iborra, F. J. & Pires Das Neves, R. Osmotic modulation of chromatin impacts on efficiency and kinetics of cell fate modulation. Sci. Rep. 8, 1–14 (2018).

52. Guilak, F., Erickson, G. R. & Ting-Beall, H. P. The effects of osmotic stress on the viscoelastic and physical properties of articular chondrocytes. Biophys. J. 82, 720–727 (2002).

53. Welsh, T. J., Krainer, G., Espinosa, J. R., Joseph, J. A., Sridhar, A., Jahnel, M., Arter, W. E., Saar, K. L., Alberti, S., Collepardo-Guevara, R. & Knowles, T. P. J. Surface Electrostatics Govern the Emulsion Stability of Biomolecular Condensates. Nano Lett. 22, 612–621 (2022).

54. Brangwynne, C. P., Tompa, P. & Pappu, R. V. Polymer physics of intracellular phase transitions. Nat. Phys. 11, 899–904 (2015).

55. Allahverdi, A., Chen, Q., Korolev, N. & Nordenskiöld, L. Chromatin compaction under mixed salt conditions: Opposite effects of sodium and potassium ions on nucleosome array folding. Sci. Rep. 5, 8512 (2015).

56. Finnin, M. S., Donigian, J. R., Cohen, A., Richon, V. M., Rifkind, R. A., Marks, P. A., Breslow, R. & Pavletich, N. P. Structures of a histone deacetylase homologue bound to the TSA and SAHA inhibitors. Nature 401, 188–193 (1999).

57. Xia, J., Zhao, J. Z., Strom, A. R. & Brangwynne, C. P. Chromatin heterogeneity modulates nuclear condensate dynamics and phase behavior. Nat. Commun. 16, 6406 (2025).

58. Hu, X., Zhang, X., Dai, L., Zhu, J., Jia, Z., Wang, W., Zhou, C. & Ao, Y. Histone deacetylase inhibitor trichostatin A promotes the osteogenic differentiation of rat adipose-derived stem cells by altering the epigenetic modifications on Runx2 promoter in a BMP signaling-dependent manner. Stem Cells Dev. 22, 248–255 (2013).

59. Hergeth, S. P. & Schneider, R. The H1 linker histones: multifunctional proteins beyond the nucleosomal core particle. EMBO Rep. 16, 1439–1453 (2015).

60. Clark, D. J. & Kimura, T. Electrostatic mechanism of chromatin folding. J. Mol. Biol. 211, 883–896 (1990).

61. Willcockson, M. A. et al. H1 histones control the epigenetic landscape by local chromatin compaction. Nature 589, 293–298 (2021).

62. Pedersen, S. F., Hoffmann, E. K. & Mills, J. W. The cytoskeleton and cell volume regulation. *Comp. Biochem. Physiol. Part A*, Mol. Integr. Physiol. 130, 385–399 (2001).

63. Hashemi, H. S., Fallone, S., Boily, M., Towers, A., Kilgour, R. D. & Rivaz, H. Assessment of mechanical properties of tissue in breast cancer-related lymphedema using ultrasound elastography. IEEE Trans. Ultrason. Ferroelectr. Freq. Control 66, 541–550 (2019).

64. Gill, S., Wight, T. N. & Frevert, C. W. Proteoglycans: Key regulators of pulmonary inflammation and the innate immune response to lung infection. Anat. Rec. 293, 968–981 (2010).

65. Distler, J. H. W., Györfi, A. H., Ramanujam, M., Whitfield, M. L., Königshoff, M. & Lafyatis, R. Shared and distinct mechanisms of fibrosis. Nat. Rev. Rheumatol. 15, 705–730 (2019).

66. Fan, W. et al. Matrix viscoelasticity promotes liver cancer progression in the pre-cirrhotic liver. Nature 626, 635–642 (2024).

67. Soares Da Costa, D., Reis, R. L. & Pashkuleva, I. Sulfation of glycosaminoglycans and its implications in human health and disorders. Annu. Rev. Biomed. Eng. 19, 1–26 (2017).

68. Agbani, E. O. & Poole, A. W. Procoagulant platelets: generation, function, and therapeutic targeting in thrombosis. Blood 130, 2171–2179 (2017).

69. Reddy, E. C. & Rand, M. L. Procoagulant Phosphatidylserine-Exposing Platelets in vitro and in vivo. Front. Cardiovasc. Med. 7, (2020).

70. Strebel, A., Lickert, S., Klamroth, R., Vogel, V. & Pennacchio, F. A. independently of thrombin generation. Blood Vessel. Thromb. Hemost. 3, 100132 (2026).

71. Singh, S., Dodt, J., Volkers, P., Hethershaw, E., Philippou, H., Ivaskevicius, V., Imhof, D., Oldenburg, J. & Biswas, A. Structure functional insights into calcium binding during the activation of coagulation factor XIII A. Sci. Rep. 9, 1–18 (2019).

72. Lueckgen, A., Garske, D. S., Ellinghaus, A., Mooney, D. J., Duda, G. N. & Cipitria, A. Enzymatically-degradable alginate hydrogels promote cell spreading and in vivo tissue infiltration. Biomaterials 217, 119294 (2019).

73. Orthaber, D., Bergmann, A. & Glatter, O. SAXS experiments on absolute scale with Kratky systems using water as a secondary standard. J. Appl. Crystallogr. 33, 218–225 (2000).

74. Patro, R., Duggal, G., Love, M. I., Irizarry, R. A. & Kingsford, C. Salmon provides fast and bias-aware quantification of transcript expression. Nat. Methods 14, 417–419 (2017).

75. Frankish, A. et al. Gencode 2021. Nucleic Acids Res. 49, D916–D923 (2021).

76. Love, M. I., Soneson, C., Hickey, P. F., Johnson, L. K., Tessa Pierce, N., Shepherd, L., Morgan, M. & Patro, R. Tximeta: Reference sequence checksums for provenance identification in RNA-seq. PLoS Comput. Biol. 16, 1–13 (2020).

77. Love, M. I., Huber, W. & Anders, S. Moderated estimation of fold change and dispersion for RNA-seq data with DESeq2. Genome Biol. 15, 1–21 (2014).

78. Maechler, M., Rousseeuw, P., Struyf, A., Hubert, M. & Hornik, K. cluster: Cluster Analysis Basics and Extensions. https://cran.r-project.org/package=cluster (2022).

79. Wu, T., Hu, E., Xu, S., Chen, M., Guo, P., Dai, Z., Feng, T., Zhou, L., Tang, W., Zhan, L., Fu, X., Liu, S., Bo, X. & Yu, G. clusterProfiler 4.0: A universal enrichment tool for interpreting omics data. Innovation 2, 100141 (2021).

80. Yang, X., Dargaville, B. L. & Hutmacher, D. W. Elucidating the Molecular Mechanisms for the Interaction of Water with Polyethylene Glycol-Based Hydrogels: Influence of Ionic Strength and Gel Network Structure. Polymers (Basel). 13, 845 (2021).

81. Posbeyikian, A., Tubert, E., Bacigalupe, A., Escobar, M. M., Santagapita, P. R., Amodeo, G. & Perullini, M. Evaluation of calcium alginate bead formation kinetics: An integrated analysis through light microscopy, rheology and microstructural SAXS. Carbohydr. Polym. 269, 1–10 (2021).

82. Yuguchi, Y., Urakawa, H., Kajiwara, K., Draget, K. I. & Stokke, B. T. Small-angle X-ray scattering and rheological characterization of alginate gels. 2. Time-resolved studies on ionotropic gels. J. Mol. Struct. 554, 21–34 (2000).

83. Hashemnejad, S. M. & Kundu, S. Rheological properties and failure of alginate hydrogels with ionic and covalent crosslinks. Soft Matter 15, 7852–7862 (2019).

84. Stokke, B. T., Draget, K. I., Smidsrød, O., Yuguchi, Y., Urakawa, H. & Kajiwara, K. Small-Angle X-ray Scattering and Rheological Characterization of Alginate Gels. 1. Ca−Alginate Gels. Macromolecules 33, 1853–1863 (2000).

85. Debye, P., Anderson, H. R. & Brumberger, H. Scattering by an inhomogeneous solid. II. the correlation function and its application. J. Appl. Phys. 28, 679–683 (1957).

86. Pedersen, J. S. Analysis of small-angle scattering data from colloids and polymer solutions: Modeling and least-squares fitting. Adv. Colloid Interface Sci. 70, 171–210 (1997).

87. Glatter, O. Scattering Methods and their Application in Colloid and Interface Science. Scattering Methods and their Application in Colloid and Interface Science (Elsevier, 2018). doi:10.1016/b978-0-12-813580-8.00016-x.

